# 5-HT_1A_ and 5-HT_2B_ receptor interaction and co-clustering regulates serotonergic neuron excitability

**DOI:** 10.1101/2022.12.09.519723

**Authors:** Amina Benhadda, Célia Delhaye, Imane Moutkine, Xavier Marques, Marion Russeau, Corentin Le Magueresse, Anne Roumier, Sabine Lévi, Luc Maroteaux

**Affiliations:** Institut du Fer à Moulin, U1270 INSERM, Sorbonne Université, 17 rue du Fer à Moulin 75005, Paris, France

## Abstract

Many psychiatric diseases including depression, schizophrenia and anxiety have been associated with serotonin (5-HT) neuron dysfunction. Pacemaker-like firing of raphe 5-HT neurons was proposed to be under unique 5-HT_1A_ receptor-mediated autoinhibition. We previously showed that 5-HT_2B_ receptors were expressed by 5-HT neurons together with 5-HT_1A_ receptors. However, functional consequences on 5-HT neurons of putative interaction between these receptors are unknown. Using co-immunoprecipitation, BRET, confocal and super-resolution microscopy in hippocampal and 5-HT neurons, we present converging evidence that 5-HT_1A_ and 5-HT_2B_ receptors can form heterodimers and co-cluster at the surface of dendrites. 5-HT_2B_ receptor clusters were redistributed upon 5-HT_1A_ receptor expression supporting functional interactions between the two receptors. Furthermore, 5-HT_2B_ receptor expression prevented agonist-induced internalization of 5-HT_1A_ receptors, whereas 5-HT_1A_ receptors mimicked the clustering effect of 5-HT_2B_ receptor stimulation on its surface expression. The functional impact of this interaction *in-vivo* was assessed by recording 5-HT neuron excitability from mice lacking 5-HT_2B_ receptors in 5-HT neurons. Upon 5-HT_1A_ receptor stimulation, the firing activity of 5-HT neurons was increased in the absence of 5-HT_2B_ receptors and decreased in their presence through regulation of SK channels, thus demonstrating functional output of this interaction in controlling 5-HT neuron firing activity.

## INTRODUCTION

Many psychiatric diseases including depression, schizophrenia or anxiety have been associated with serotonin (5-HT) neuron activity dysfunctions (Mann, 1999; Arango et al., 2003; Marazziti, 2017). Dorsal raphe serotonin neurons (DR) and their long-range projections exert control over many physiological functions such as emotion, sleep, and locomotion. With the exception of 5-HT_3_ receptors (5-HT_3_-Rs), which are ion channels, all 5-HT-Rs are G-protein coupled receptors (GPCR) that differ according to their G-protein coupling.

The 5-HT_1_-Rs subtypes are coupled to the Gαi/o protein and their activation leads to a decrease in intracellular cAMP (Lin et al., 2002). 5-HT_1A_-Rs are expressed in the central nervous system as an autoreceptor (expressed by 5-HT neurons), and as a heteroreceptor in neurons from several brain structures including hippocampus and prefrontal cortex (Garcia-Garcia et al., 2014; Andrade et al., 2015; Albert and Vahid-Ansari, 2019). In 5-HT neurons, 5-HT_1A_-R expression is located to the somatodendritic compartment, and 5-HT_1B_-R (a 5-HT_1A_-R close relative) expression is restricted to synaptic projections (McDevitt and Neumaier, 2011). Through the Gβγ subunit, 5-HT_1A_-R activation can induce the opening of potassium channels (G protein-coupled inwardly-rectifying potassium channels or GIRKs) that hyperpolarize 5-HT neurons and inhibit their firing (Heusler et al., 2005; Llamosas et al., 2015; Montalbano et al., 2015; Courtney and Ford, 2016). The serotonergic tone of the DR neurons has thus been considered to be exclusively regulated by 5-HT_1A_-Rs expressed on their dendritic surface (autoreceptors): activity-dependent release of 5-HT within serotonergic raphe nuclei activates these 5-HT_1A_ autoreceptors, opens GIRK channels, hyperpolarizes 5-HT neurons, and thereby inhibits cell firing, thus completing an autoinhibitory feedback loop (Montalbano et al., 2015). However, several accumulating evidence indicate alternative roles for 5-HT_1A_-R in mediating autoinhibition in dorsal raphe beyond this homeostatic control of firing rate (Andrade et al., 2015). 5-HT_1A_ autoreceptors are subject to homologous and heterologous desensitization, which are the result of receptor internalization (Riad et al., 2001) and are thought to participate into chronic serotonin-specific reuptake inhibitors (SSRIs) antidepressant effects. However, there is also evidence for persistent 5-HT_1A_-R-dependent feedback inhibition of 5-HT neurons after sustained exposure to high extracellular 5-HT as upon exposure to SSRIs (Soiza-Reilly et al., 2015) questioning its exact role. Small-conductance Ca^2+^-activated potassium (SK) channels are responsible for the medium afterhyperpolarisation (mAHP) that follows action potentials in neurons (Kirby et al., 2003). Ca^2+^ regulation of SK channels was shown to control neuron firing by acting at amplitude and duration of mAHPs and to switches pacemaker-like firing pattern to burst firing (repeated action potentials separated by a small interval). SK channels are thus able to tune the excitability of monoamine neurons (Ji et al., 2009). Apamin, a neurotoxin from bee venom, which is a potent antagonist of SK channels and in particular of SK2 and SK3 subtypes, has been shown to increase 5-HT neuron burst firing in DR and extracellular 5-HT levels (Crespi, 2009) via a direct action (Rouchet et al., 2008). Nevertheless, regulation of the firing pattern of 5-HT neurons by autoreceptors is still poorly understood (Andrade et al., 2015).

The 5-HT_2_-Rs subtypes (5-HT_2A, 2B, 2C_) are coupled to the Gαq/11 protein that activates phospholipase C (PLC) and increases inositol triphosphate (IP3) and Ca^2+^ intracellular concentration. A loss of function of *HTR2B*, the gene encoding 5-HT_2B_-Rs, is associated with impulsivity, a higher risk of developing depression and suicidal behavior (Bevilacqua et al., 2010; Tikkanen et al., 2015). Using behavioral and biochemical experiments in mice, Diaz et al. (2012) showed that 5-HT_2B_-Rs are expressed with 5-HT_1A_-Rs in serotonergic neurons as later validated by RNAseq (Niederkofler et al., 2016; Okaty et al., 2020) (see https://bokaty.shinyapps.io/DR_Pet1_neuron_scRNAseq_DB/). The acute or chronic activation of 5-HT_2B_-Rs by the preferential agonist BW723C86 (BW) was able to mimic SSRI effects in mice (Diaz et al., 2012). In mice lacking the 5-HT_2B_-R specifically in 5-HT neurons (cKO^5-HT^ mice), behavioral and molecular responses to SSRIs are abolished. Moreover, in WT mice, the activation of 5-HT_2B_-Rs by BW was able to counteract the effects of 5-HT_1A_ autoreceptor activation, such as the decrease in 5-HT neurons firing rate and in body temperature (Belmer et al., 2018). Previous work by our team revealed that the expression of 5-HT_2B_-Rs in hippocampal neurons was restricted to the somatodendritic compartment, where they formed clusters (Benhadda et al., 2021). Although 5-HT_2_-Rs have been shown to crosstalk (Kidd et al., 1991), and 5-HT_2B_-Rs are expressed by serotonergic neurons together with 5-HT_1A_-Rs (Diaz et al., 2012), putative functional interactions between these receptors and their consequence to serotonergic tone regulation are not yet documented.

The aim of this work was thus to test the hypothesis that 5-HT_2B_-Rs interact with 5-HT_1A_-Rs and that this interaction affects their respective intracellular distribution and thus their physiological functions on 5-HT neurons. By studying their expression in different cell types (COS-7 cells, primary cultures of hippocampal neurons, and *in-vivo* raphe 5-HT neurons) using various pharmacological, electrophysiological, and high-resolution imaging techniques, we show that 5-HT_2B_-Rs can form dimers with 5-HT_1A_-Rs. This interaction has no apparent effect on their second messenger signalization and their respective ligand binding properties. However, we found that this interaction alters the cell-surface expression of these receptors (clustering, redistribution) during responses to specific ligands and thus their signaling efficacy that ultimately controls the excitability of 5-HT neurons *in-vivo* through SK and GIRK channels regulation.

## RESULTS

### Coexpression of 5-HT_2B_-Rs with 5-HT_1A_-Rs reveals direct interactions

We previously showed that 5-HT_2B_-Rs and 5-HT_1A_-Rs were coexpressed in 5-HT neurons (Diaz et al., 2012) and have opposite action on their activity (Belmer et al., 2018) that may result from reciprocal interactions. To assess putative 5-HT_2B_-R and 5-HT_1A_-R interactions, we first performed co-immunoprecipitation experiments in COS-7 cells co-transfected with plasmids encoding HA-tagged 5-HT_2B_-Rs and/or myc-tagged 5-HT_1A_-Rs. Extracted proteins were immunoprecipitated using anti-HA beads followed by Western blotting revealing co-immunoprecipitation between 5-HT_1A_-Rs and 5-HT_2B_-Rs (**Fig. 1A**). We completed these data by performing BRET experiments to assess putative close interactions between 5-HT_1A_-Rs and 5-HT_2B_-Rs. This experiment relies on the natural capacity of Renilla luciferase, used here as BRET donor, to emit energy in presence of its substrate, coelenterazine or to transfer it to an acceptor. Luciferase coding region and yellow fluorescent protein (YFP, here, BRET acceptor) were fused to the C-terminal coding region of 5-HT_1A_-Rs and 5-HT_2B_-Rs. Constant amounts of BRET donor plasmids and increasing amounts of BRET acceptor plasmids were transfected in COS-7 cells. A hyperbolic curve confirms the specific interactions between the BRET donor with the acceptor and thus, the close proximity between the two receptors studied. Hyperbolic curve was obtained when 5-HT_2B_-Rs and 5-HT_1A_-Rs were expressed together supporting that they can form heterodimers in intact cells (**Fig. 1B**). As control, no BRET signal was obtained when 5-HT_2B_-Rs were tested for self-association, as shown in a previous study indicating that these receptors were unable to form homodimers (Moutkine et al., 2017). Thus, co-immunoprecipitation and BRET experiments revealed that 5-HT_1A_-Rs and 5-HT_2B_-Rs can interact closely and may form heteromers.

### Total receptor expression and second messenger coupling are not affected by 5-HT2B-R /5-HT1A-R coexpression

We assessed if coexpression of 5-HT_1A_-Rs and 5-HT_2B_-Rs affect their respective intracellular signaling pathway, and pharmacological properties. Radioligand binding competition by increasing concentrations of BW or NAN of tritiated mesulergine or tritiated 8-OH-DPAT, for 5-HT_2B_-R and 5-HT_1A_-R, respectively, was performed on total membrane proteins from transfected COS-7 cells expressing 5-HT_2B_-Rs with or without 5-HT_1A_-Rs. No difference was found in total expression of the receptors, which is represented by the Bmax, that reflects the number of available receptors, nor in binding affinity to their respective ligand (Ki) in coexpressing compared to single receptor expressing cells (**Fig. 2A-B**). Using HTRF-based assays, we measured the intracellular inositolphosphate (IP) amount produced by agonist stimulation of 5-HT_2B_-Rs. Dose-response curve of 5-HT_2B_-R-mediated production of IP in response to BW in cells expressing 5-HT_2B_-Rs alone or in combination with 5-HT_1A_-Rs were analyzed using the operational model to determine the Gαq coupling efficiency of single receptor and heterodimer. The dose response curves showed no significant difference in the EC50 between the 2 conditions (**Fig. 2C**). These results suggest that 5-HT_1A_-R and 5-HT_2B_-R coexpression modifies neither their expression nor the 5-HT_2B_-R coupling efficiency.

**Figure 1:**
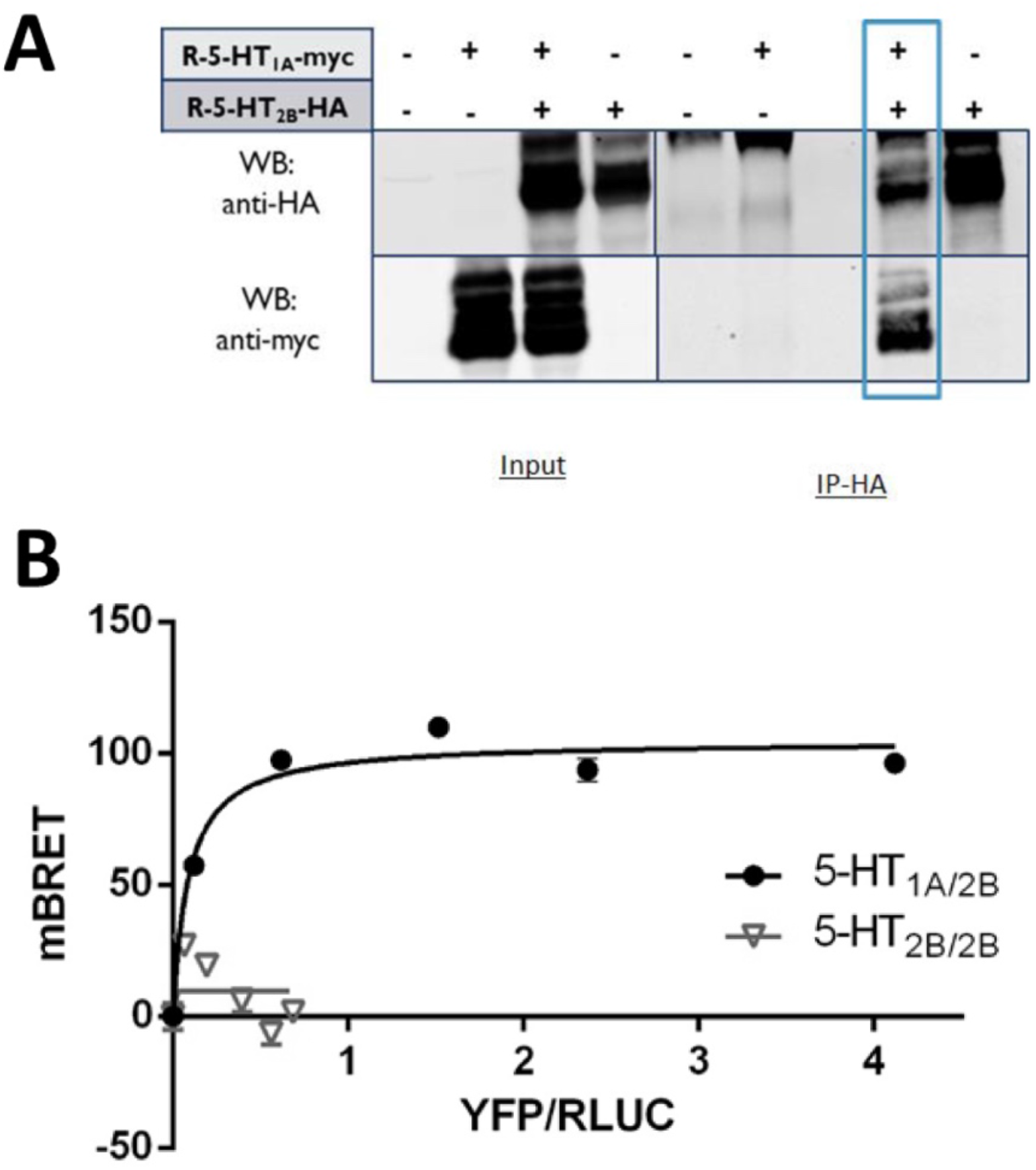
Physical proximity of 5-HT_2B_-Rs and 5-HT_1A_-Rs in transfected cells. (**A**) Co-immunoprecipitation of 5-HT_1A_-Rs coexpressed with 5-HT_2B_-Rs in COS-7 cells. Proteins from COS-7 cells co-transfected with plasmids coding for HA-tagged 5-HT_2B_ (R-5-HT_2B_-HA) and myc-tagged 5-HT_1A_ (R-5-HT_1A_-myc) receptors were immunoprecipitated and analyzed by Western blotting. Blot of input of proteins revealed by anti-myc antiserum (WB anti-myc) or by anti-HA antiserum (WB anti-HA) shows the presence of the proteins in each condition (Input; left panel). Immunoprecipitations using anti-HA beads (IP-HA; right panel) revealed a co-immunoprecipitation between 5-HT_1A_-Rs and 5-HT_2B_-Rs. (**B**) BRET proximity assays for 5-HT_1A_-Rs and 5-HT_2B_-Rs. COS-7 cells were co-transfected with a constant amount of plasmids coding for RLuc-5-HT_2B_-R (BRET donor) and increasing amount of plasmid encoding YFP-tagged 5-HT_1A_-R (BRET acceptors) or YFP-tagged 5-HT_2B_-R as control unable to form dimers (Moutkine et al., 2017). The hyperbolic curve attests close interaction between 5-HT_1A_-YFP and 5-HT_2B_-RLuc with a physical distance <10 nm (representative experiment of n > 3 independent experiments).

**Figure 2:**
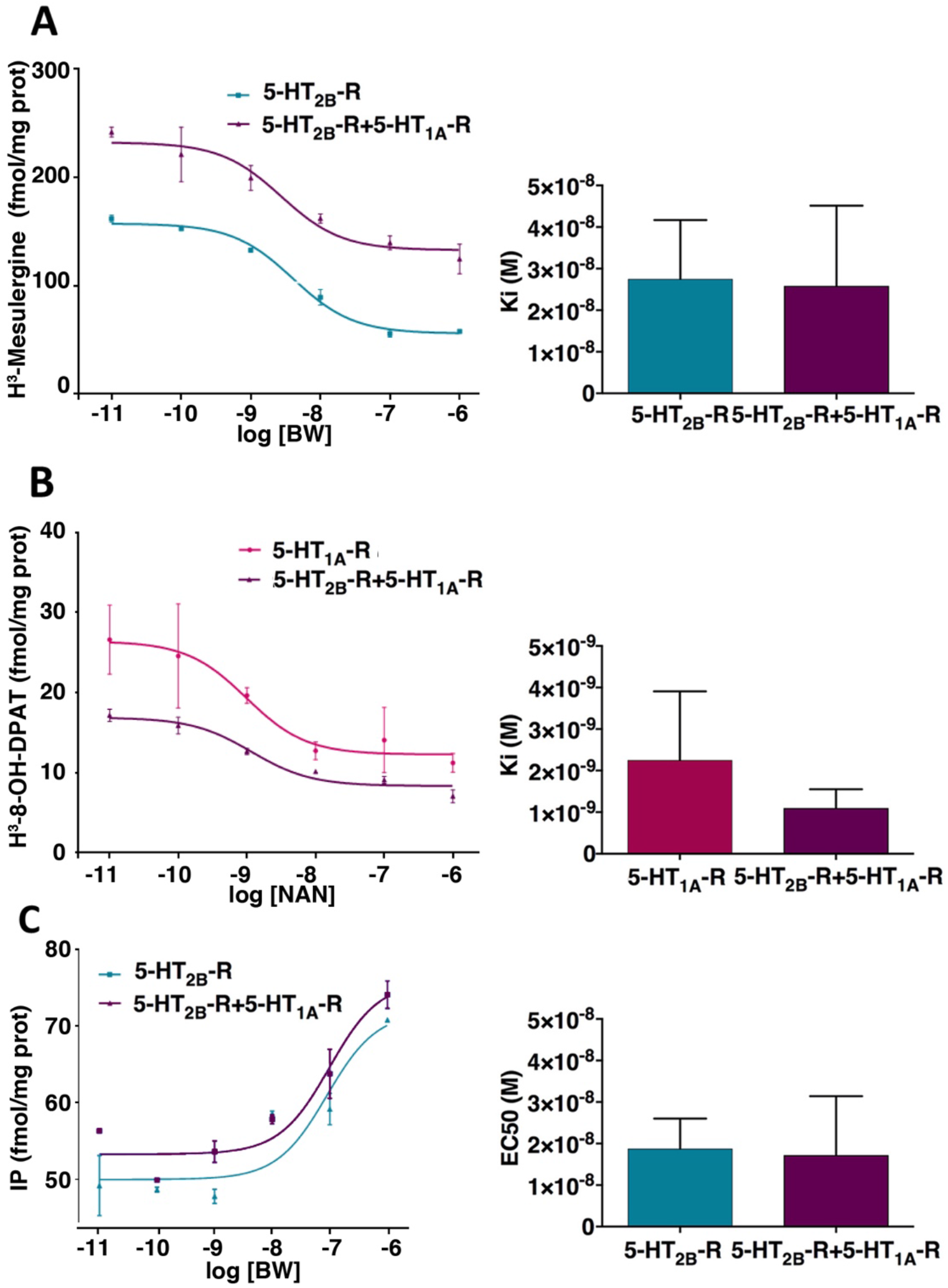
Receptor expression and second messenger signaling are not affected by 5-HT_2B_-R /5-HT_1A_-R dimerization. Radioligand binding competition by BW or NAN of tritiated-mesulergine **(A**) or 8-OH-DPAT **(B**) performed on total membrane proteins from transfected COS-7 cells expressing 5-HT_2B_-Rs with or without 5-HT_1A_-Rs. No effect was found in the coexpression condition regarding the total receptor expression (Bmax) or their binding affinity to their respective ligand (Ki) (Right). Data were analyzed using the Mann-Whitney unpaired t-test. Representative curves of n = 3 independent experiments, each performed in duplicate. (**C**) Impact of coexpression on 5-HT_2B_-R- and 5-HT_1A_-R-operated signal transduction. The quantification of inositol phosphate was performed in COS-7 cells expressing 5-HT_2B_-Rs with or without 5-HT_1A_-Rs and stimulated with increasing concentrations of the 5-HT_2B_-R agonist BW. Note the absence of effect, in the coexpression condition, on IP production compared to the single 5-HT_2B_-R expression condition. Data were statistically analyzed by Mann-Whitney unpaired t-test. Representative curves of n = 3 independent experiments, each performed in duplicate.

### 5-HT_2B_-Rs/5-HT_1A_-Rs coexpression decreases 5-HT_2B_-R surface expression

As the total expression level and the coupling properties of the receptors were not modified by their coexpression, we investigated putative changes in their cellular distribution. First, we performed surface binding assays on intact cells in order to evaluate their plasma membrane density. Using similar pharmacological protocol on non-permeabilized living cells, we observed that coexpression of the two receptors did not modify affinity but decreased the amount of 5-HT_2B_-R surface expression as evaluated by their Bmax; compared to 5-HT_2B_-Rs expressed alone, Bmax was significantly lower (by 60%) in coexpression condition with 5-HT_1A_-Rs (**Fig. 3A**) but no difference was found in 5-HT_1A_-R plasma membrane expression (not illustrated). To confirm this result, we performed surface biotinylation assay, which is a direct method to evaluate protein surface expression. To overcome fluctuations of total protein expression in different transfection, results were analyzed by the ratio of the surface/total protein expression for each receptor. The results showed that coexpression of the two receptors significantly decreased 5-HT_2B_-R surface by 31% but not 5-HT_1A_-R surface expression (**Fig. 3B-C**). Together, these data indicate that the 5-HT_2B_-R surface expression is reduced by coexpression of 5-HT_1A_-Rs, supporting functional interactions between the two receptors.

**Figure 3:**
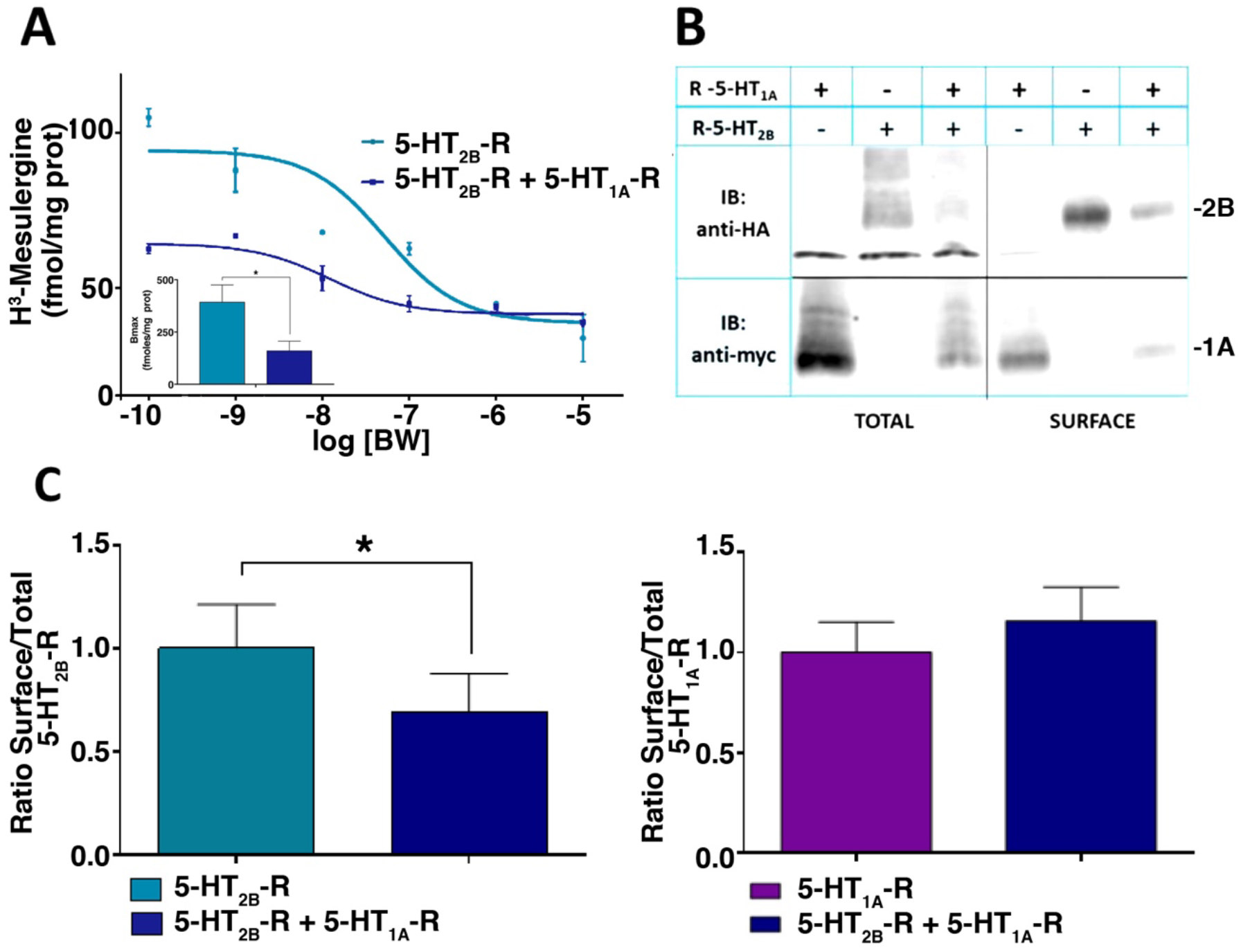
5-HT_2B_-R /5-HT_1A_-R co-expression decreases 5-HT_2B_-R surface expression. **(A**). Radioligand binding competition by BW of tritiated mesulergine performed on surface proteins (intact cells) from transfected COS-7 cells expressing 5-HT_2B_-Rs with or without 5-HT_1A_-Rs. Co-expression revealed a decreased surface expression of the 5-HT_2B_-R illustrated by a significant decrease in Bmax (p=0.037; inset). Data were analyzed using unpaired t-test with Welch’s correction; n = 3 independent experiments, each performed in duplicate. (**B-C**). Surface biotinylation performed on COS-7 cells expressing 5-HT_1A_-Rs and/or 5-HT_2B_-Rs. (**B**) Analysis by Western blotting. (**C**). Ratio of the surface/total protein expression for each receptor analyzed in conditions of simple transfection or in co-expression experiments. Co-expression of the two receptors confirmed the decrease in 5-HT_2B_-R (unpaired t-test p=0.0474, n = 5) but not 5-HT_1A_-R surface expression (unpaired t-test p=0.4596, n=5). Representative curves of n = 5 independent experiments, each performed in duplicate. Band intensity of each condition was quantified and represented graphically (*p<0,05).

### Neuronal distribution of 5-HT_2B_-Rs in somatodendritic compartment as 5-HT_1A_-Rs

We previously showed that 5-HT_2B_-R expression was restricted to somatodendritic compartment of hippocampal cultured neurons (Benhadda et al., 2021), and that 5-HT_2B_-Rs can positively regulate 5-HT neuronal activity in an opposite manner to 5-HT_1A_-Rs (Belmer et al., 2018). Independently, 5-HT_1A_-Rs were shown to be expressed in the somatodendritic compartment of either 5-HT or hippocampal neurons *in-vivo* (Kia et al., 1996; Riad et al., 2000) and in cultures (Renner et al., 2012). However, even if 5-HT_1A_-Rs and 5-HT_2B_-Rs are both endogenously expressed by 5-HT neurons and hippocampal neurons, the lack of selective antibodies against these receptors makes studies on endogenous receptors impossible. We thus transfected hippocampal neuron cultures with plasmids encoding HA-tagged-5-HT_2B_-Rs and myc-tagged-5-HT_1A_-Rs to study their cellular and subcellular distribution. Although these two receptors were expressed in the somatodendritic compartment, 5-HT_1A_-Rs and 5-HT_2B_-Rs displayed different expression patterns: 5-HT_2B_-Rs were arranged in large clusters, whereas 5-HT_1A_-Rs were detected as much smaller clusters at the surface of dendrites of hippocampal neurons (**Fig. 4A**). We confirmed in hippocampal cultured neurons that the coexpression of 5-HT_1A_-Rs decreased the surface expression of 5-HT_2B_-Rs by 40% compared to its total expression (**Fig. 4A**), as previously observed in COS cells (**Fig. 3C**). Next, we developed pAAV-EF1A-DIO-WPRE-pA viruses for stereotaxic injection into the B7 raphe nuclei of *Pet1-Cre*^*+/0*^ mice in order to express Flag-tagged-5-HT_1A_-Rs and HA-tagged-5-HT_2B_-Rs specifically in DR 5-HT neurons. By this mean, we showed somatodendritic colocalization of both receptors in 5-HT neurons *in-vivo* (**Fig. 4B**). Together, these data suggest that 5-HT_2B_-Rs and 5-HT_1A_-Rs are in close proximity and may functionally interact in neurons.

**Figure 4:**
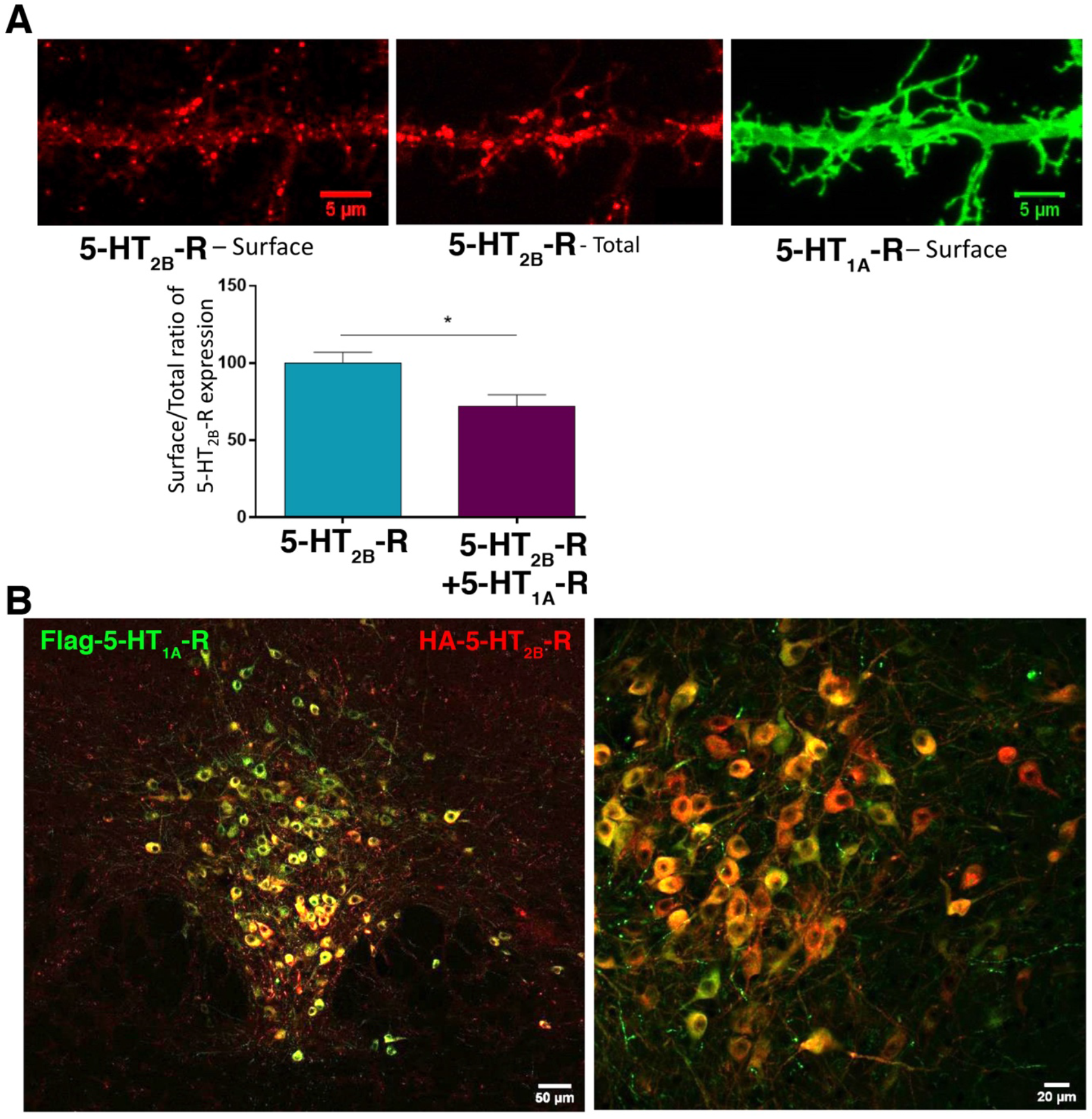
Co-expression of 5-HT_2B_-Rs with 5-HT_1A_-Rs in hippocampal and raphe neurons. (**A**). 5-HT_2B_-R localization by HA-tag immunostaining revealed receptor clustering (red) at the dendritic surface of hippocampal neurons, whereas the 5-HT_1A_-R localization studied using myc-tag immunostaining reported a more even distribution in small clusters (green) of 5-HT_1A_-R. Furthermore, a staining in the absence of detergent (surface expression) or in the presence of detergent (total expression) indicated that co-expressing 5-HT_1A_-Rs with 5-HT_2B_-Rs decreased the surface/total ratio of 5-HT_2B_-Rs. (p=0.011, as assessed by unpaired t-test; n=3 independent experiments; Scale bars, 5 µm). **(B**). Mouse brains were examined to assess the distribution of adeno-associated virus-encoded DIO Flag-tagged 5-HT_1A_-Rs and HA-tagged 5-HT_2B_-Rs injected specifically in B7 raphe 5-HT neurons of *Pet1-Cre*^*+/0*^ mice. Images of 5-HT neurons labeled with Flag (green) and HA (red) antibodies is shown at increasing magnifications (from left to right) and reveals colocalization of both receptors in the somatodendritic compartment. Scale bars, 50 and 20 µm, as indicated.

### Stimulation of 5-HT_2B_-Rs increases its surface clustering and maintains expression of 5-HT_1A_-Rs at cell surface

In primary cultures of hippocampal neurons, we studied the subcellular distribution of both receptors using confocal microscopy. We quantified the number and size of 5-HT_2B_-R clusters and the density of molecules per cluster (integrated intensity) with or without 5-HT_1A_-R coexpression and following selective stimulation with the 5-HT_2B_-R agonist BW (1 µM). BW stimulation of 5-HT_2B_-Rs for 20 min decreased the number of 5-HT_2B_-R clusters (51%), increased the size (97%) and density of receptor per cluster (64%) (**Fig. 5A**). In the absence of stimulation, coexpression of 5-HT_1A_-Rs decreased the number of 5-HT_2B_-R clusters by 36% and increased their area by 73%, but did not alter their density (**Fig. 5A**), and selective stimulation by BW did not significantly alter these parameters. Selective stimulation of 5-HT_1A_-R by 8-OH-DPAT (1 µM) led to a reduction of 5-HT_1A_-R expression at cell-surface, as shown by the decrease in surface 5-HT_1A_-R fluorescence intensity over time (41%), which suggests an internalization process (**Fig. 5B**). Coexpression with 5-HT_2B_-Rs resulted in a 34% increase in 5-HT_1A_-R expression at cell-surface, which was maintained upon 8-OH-DPAT stimulation (**Fig. 5B**). This result suggests that the presence of 5-HT_2B_-Rs increases 5-HT_1A_-R clustering and prevents its 8-OH-DPAT-induced internalization thereby confirming their interaction. Clusters are small objects, and the 200-nm diffraction limit of confocal microscopy does not allow isolation of closely spaced clusters (**Fig. 6A**). Using Stochastic Optical Reconstruction Microscopy (STORM) technique with a resolution of 20 nm, we confirmed our initial observation: coexpression, without or with 5-HT_2B_-R stimulation by BW (1 µM), significantly decreased the 5-HT_2B_-R cluster number (52% and 55%, respectively) and increased their area (292% and 150%, respectively) (**Fig. 6B-C**). Thanks to its better resolution compared to confocal microscopy, this technique revealed a small but significant potentiation by 2-fold of the effects of 5-HT_1A_-R coexpression on 5-HT_2B_-R cluster number upon BW stimulation of 5-HT_2B_-Rs (**Fig. 6B-C**). Together, these results confirmed that the coexpression of 5-HT_1A_-Rs could mimic some of the effect of 5-HT_2B_-R stimulation by BW on its surface expression.

**Figure 5:**
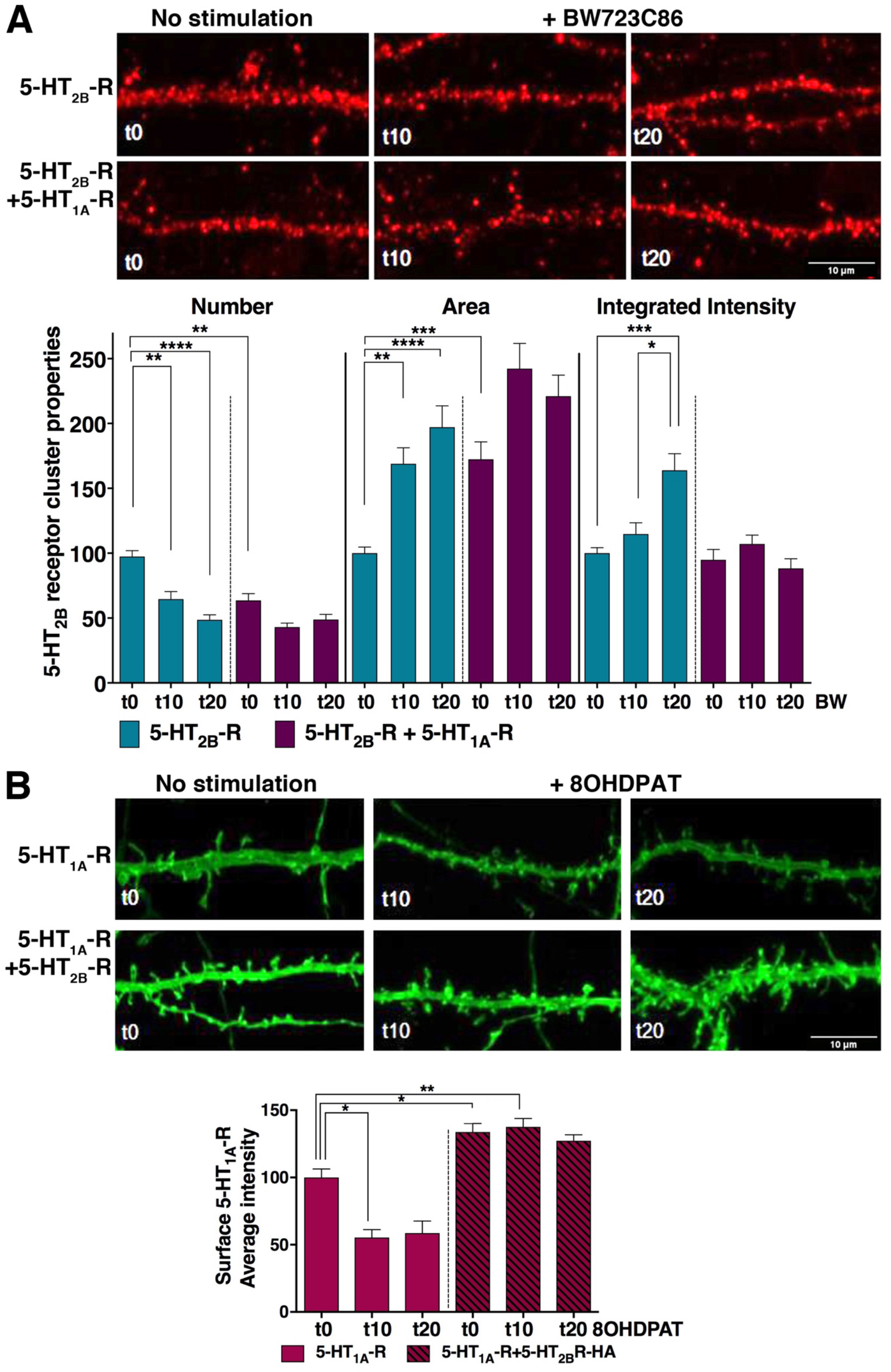
Co-expression of 5-HT_2B_-Rs/5-HT_1A_-Rs increases 5-HT_2B_-R membrane clustering and maintains 5-HT_1A_-R surface expression upon orthologous stimulation. **(A**). Top, 5-HT_2B_-R clustering at the dendritic surface 0, 10 or 20 min after BW stimulation in the absence (top) or presence (bottom) of 5-HT_1A_-Rs. Scale bars, 10 µm. Bottom, Quantification of 5-HT_2B_-R surface clustering following agonist stimulation in the presence (or not) of 5-HT_1A_-Rs. Upon 5-HT_2B_-R stimulation by BW (1 µM), the number of 5-HT_2B_-R clusters decreased in the absence of 5-HT_1A_-Rs (p=0.0026 t0 vs. t10 and p <0.0001 t0 vs. t20), as revealed by Kruskal-Wallis test, P<0.0001, followed by Dunn’s multiple comparisons test; in the presence of 5-HT_1A_-Rs, the 5-HT_2B_-R cluster number decreased in the absence of BW stimulation and was unaffected by BW stimulation, p=0.004 t0 5-HT_2B_-R vs. t0 5-HT_1A_-R. The BW stimulation of 5-HT_2B_-R alone increased the cluster size (area), p=0.0016 t0 vs. t10 and p <0.0001 t0 vs. t20; in the presence of 5-HT_1A_-Rs, the 5-HT_2B_-R cluster area increased but was unaffected by BW stimulation, p=0.0008 t0 5-HT_2B_-R vs. t0 5-HT_1A_-R. The BW stimulation slightly increased the molecular density of clusters (integrated intensity) of 5-HT_2B_-R clusters from 0 to 20 min in the absence of 5-HT_1A_-Rs, p=0.00007 t0 vs. t20; in the presence of 5-HT_1A_-Rs, the 5-HT_2B_-R cluster integrated intensity was unaffected without or with BW stimulation. Number of cells = 19-38 from 3-4 experiments. (**B**). Top, 5-HT_1A_-R expression at the dendritic surface 0, 10 or 20 min after 8-OH-DPAT stimulation in the absence (top) or presence (bottom) of 5-HT_2B_-R. Scale bars, 10 µm. Bottom, Quantification of 5-HT_1A_-R surface expression following agonist stimulation in the presence (or not) of 5-HT_2B_-Rs. Upon 5-HT_1A_-R stimulation by 8-OH-DPAT (1 µM), 5-HT_1A_-R expression at the neuronal surface decreased from 0 to 10 min in the absence of 5-HT_2B_-Rs compared to t0 as revealed by Kruskal-Wallis test, P<0.0001, followed by Dunn’s multiple comparisons test, p=0.0115 t0 vs. t10; in the presence of 5-HT_2B_-Rs, the 5-HT_1A_-R expression increased but was unaffected by 8-OH-DPAT stimulation, p=0.0172 t0 5-HT_2B_-R vs. t0 5-HT_1A_-R+5-HT_2B_-R. Number of cells = 18-43 from 3 experiments. In all graphs, data, expressed as percentage of their respective values at t0. ****p<0.0001, ***p<0.001, **p < 0.01, *p < 0.05.

**Figure 6:**
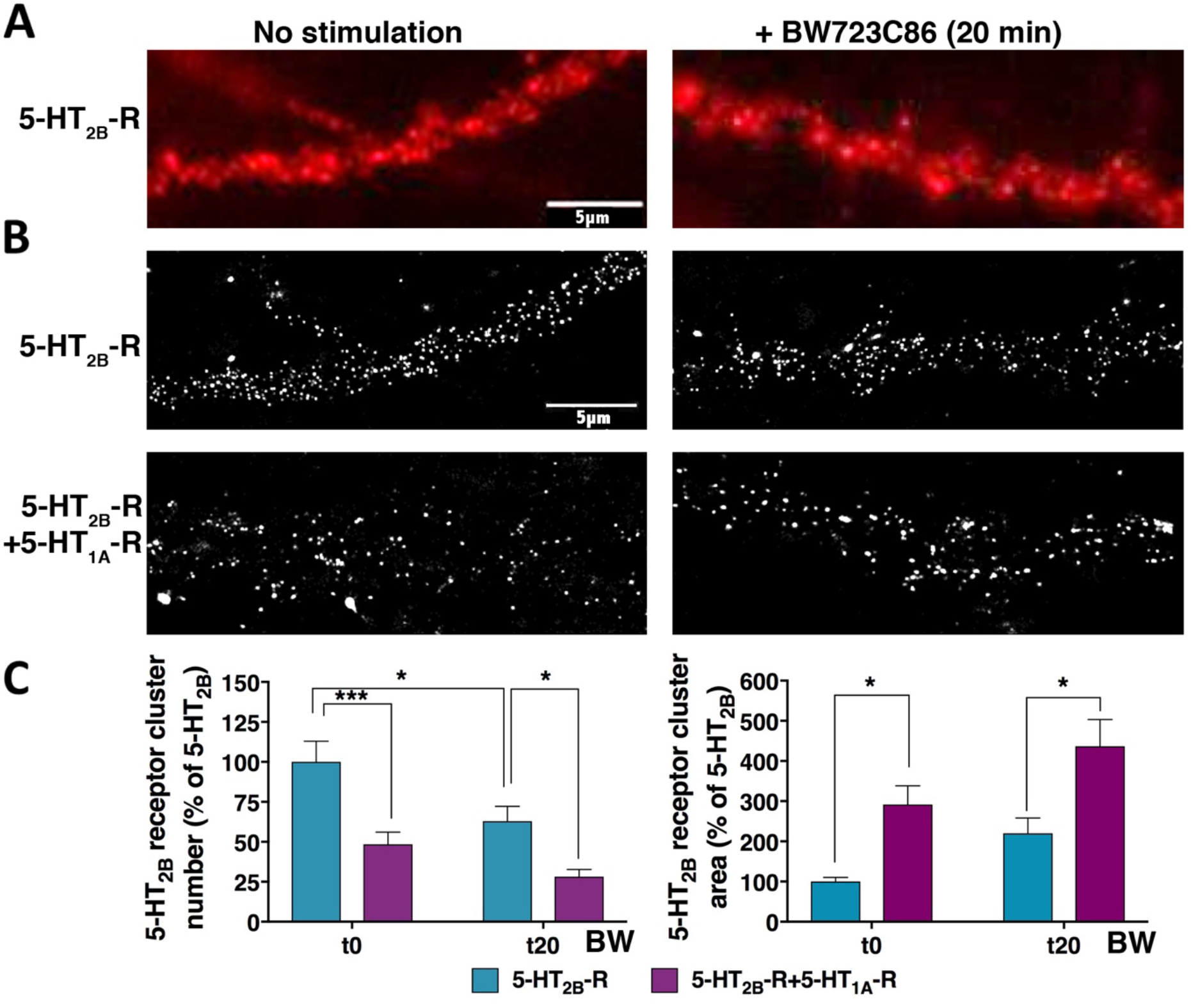
Super-resolution microscopy validated that co-expression of 5-HT_2B_-Rs with 5-HT_1A_-Rs increases 5-HT_2B_-R membrane clustering as upon BW agonist stimulation. **(A**). Images of 5-HT_2B_-R membrane clusters acquired by epifluorescence microscopy. Note that clusters are not well-defined and diffuse compared to the lower images acquired at super resolution. Scale bar, 5 µm. (**B**). STORM images of 5-HT_2B_-Rs obtained from the dendritic regions shown in A before (left panel) or 20 min (right panel) after BW stimulation in the absence (top) or presence (bottom) of 5-HT_1A_-Rs. Scale bar, 5 µm. (**C**). Quantification of 5-HT_2B_-R clustering following agonist stimulation in the presence or not of 5-HT_1A_-Rs. The number of 5-HT_2B_-R clusters was significantly decreased upon stimulation by BW (1 µM) or by co-expression with 5-HT_1A_-Rs. 2-way ANOVA revealed main effects of agonist treatment F_1, 70_ = 11.15, P=0.0014 and receptor expression F_1, 70_ = 25.31, P<0.0001; t0 5-HT_2B_-R vs. t0 5-HT_2B_-R+5-HT_1A_-R p= 0.0004; t20 5-HT_2B_-R vs. t20 5-HT_2B_-R+5-HT_1A_-R p= 0.028; t0 vs. t20 5-HT_2B_-R, p=0.0271 by Tukey’s multiple comparisons test. The area of 5-HT_2B_-R clusters was significantly increased upon stimulation by BW for 20 min (t20) or by co-expression with 5-HT_1A_-Rs. 2-way ANOVA revealed main effects of agonist treatment F_1, 70_ = 7.067, P=0.0097 and receptor expression F_1, 70_ = 16.78, P=0.0001; t0 5-HT_2B_-R vs. t0 5-HT_2B_-R+5-HT_1A_-R p= 0.0402; t20 5-HT_2B_-R vs. t20 5-HT_2B_-R+5-HT_1A_-R p= 0.0156 by Tukey’s multiple comparisons test. Number of cells = 16-21, from 3 experiments. Data are expressed as percentage of their respective values at t0. ****p<0.0001, ***p<0.001, **p < 0.01, *p < 0.05.

### Selective stimulation of either 5-HT_1A_-Rs or 5-HT_2B_-Rs generates crossed effects on receptor clustering at the cell surface

Even if no alteration of 5-HT_2B_-R transduction signaling was observed when this receptor was coexpressed with 5-HT_1A_-Rs in COS cells, the neuronal redistribution of the receptor upon stimulation is of interest since it suggests local changes in its function. In the coexpression condition, 5-HT_1A_-R surface expression increased by 28%, and when we selectively stimulated the 5-HT_2B_-R with BW (1 µM) for 20 min, we found a 28% decrease in the 5-HT_1A_-R surface expression compared to the unstimulated coexpression condition (**Fig. 7A-B**). Conversely, in the coexpression condition, we observed a 49% decrease in the 5-HT_2B_-R cluster number, and an increase in their size (by 99%) and integrated cluster intensity (by 89%) when 5-HT_1A_-Rs were selectively stimulated by 8-OH-DPAT (1 µM) for 20 min. These results indicate that stimulating 5-HT_1A_-R increases its effects on 5-HT_2B_-R clustering (**Fig. 7A-C**). These data suggest a cooperative association between 5-HT_2B_-Rs and 5-HT_1A_-Rs at the surface of neurons: the selective 5-HT_1A_-R stimulation increasing 5-HT_2B_-R clustering and the selective 5-HT_2B_-R stimulation decreasing 5-HT_1A_-R surface expression.

**Figure 7:**
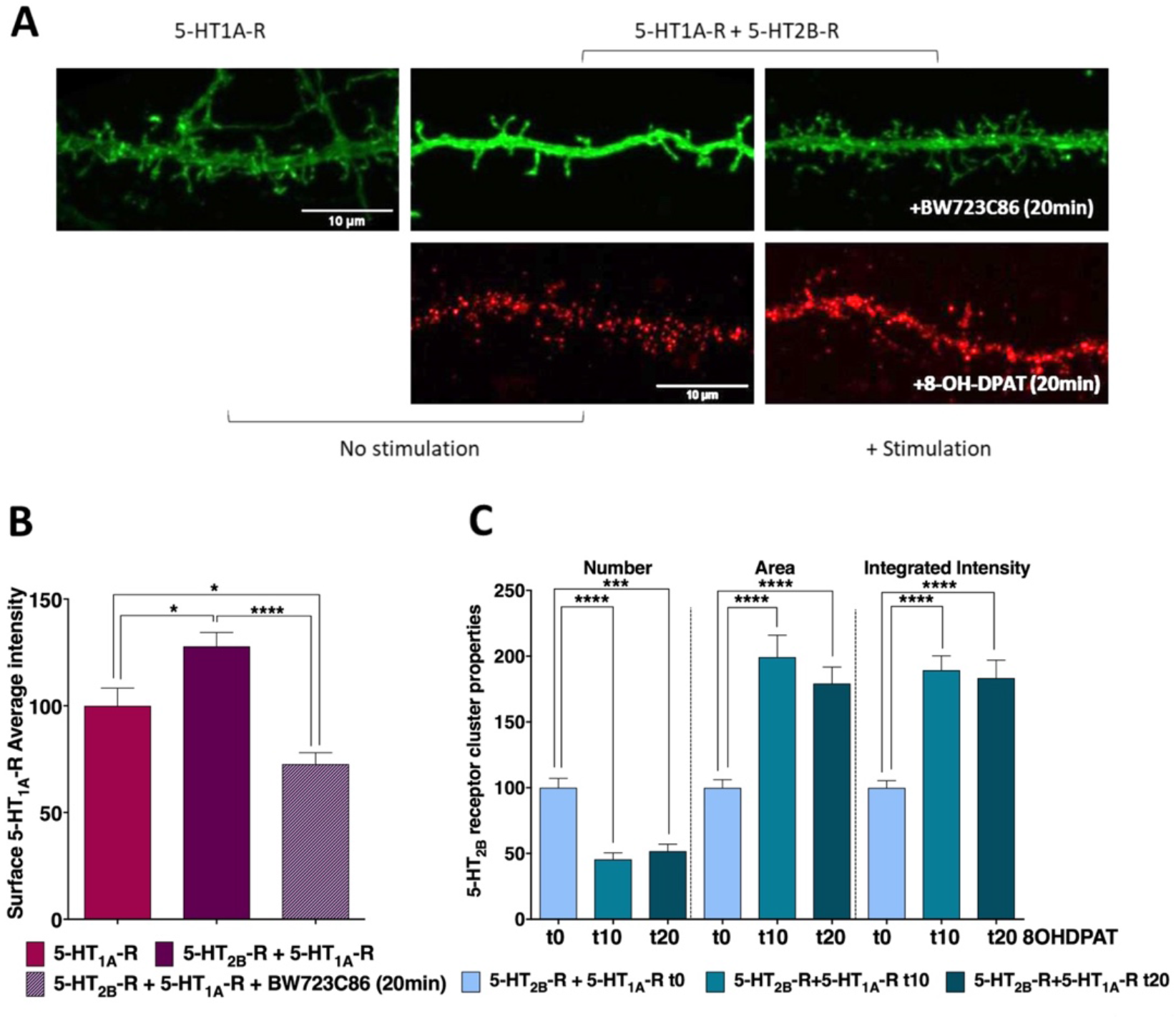
Effects of cross-stimulation of 5-HT_2B_-Rs or 5-HT_1A_-Rs on their respective surface expression. **(A**). Images of 5-HT_1A_-R and 5-HT_2B_-R surface expression acquired by confocal microscopy. Top, 5-HT_1A_-R (green) at the dendritic surface in the absence (left) or presence of 5-HT_2B_-Rs, and 0 (middle) or 20 min (right) after BW stimulation. Scale bar, 10 µm. Bottom, 5-HT_2B_-R (red) clustering at the dendritic surface 0 (left) or 20 min (right) after 8-OH-DPAT stimulation in the presence of 5-HT_1A_-Rs. Scale bar, 10 µm. (**B**) Quantification of 5-HT_1A_-R surface expression in neurons expressing 5-HT_1A_-R alone or in combination with 5-HT_2B_-Rs, with or without 5-HT_2B_-R stimulation by BW (1 µM). One-way ANOVA analysis shows a significant effect of the co-expression of 5-HT_2B_-R F_2. 43_ = 16.15, P<0.0001, with an increase in 5-HT_1A_-R surface expression, as shown by Tukey’s multiple comparison (p=0.0188 vs. non-stimulated 5-HT_1A_-R); the 5-HT_2B_-R stimulation by BW in co-expression experiment abolishes this effect (p<0.0001 vs. non-stimulated) and even reverses this effect (p=0.0463) as compared to non-stimulated neurons expressing 5-HT_1A_-R alone. Number of cells = 12-22, from 2 experiments. (**C**) Quantification of 5-HT_2B_-R clustering in neurons co-expressing 5-HT_1A_-Rs in control condition or after 10 or 20 min of 5-HT_1A_-R stimulation with 8-OH-DPAT (1 µM). Upon stimulation by 8-OH-DPAT, the number of 5-HT_2B_-R clusters decreased as revealed by Kruskal-Wallis test, P<0.0001, followed by Dunn’s multiple comparisons test; p<0.0001 t0 vs. t10 and p=0.0001 t0 vs. t20; and their area was increased, as revealed by Kruskal-Wallis test, P<0.0001, followed by Dunn’s multiple comparisons test, p<0.0001, t0 vs. t10 and p<0.0001 t0 vs. t20; their molecular density (integrated intensity) was also increased, as revealed by Kruskal-Wallis test, P<0.0001, followed by Dunn’s multiple comparisons test, p<0.0001 t0 vs. t10 and p<0.0001 t0 vs. t20. Data are expressed as percentage of their respective values at t0. Number of cells = 11-24 from 3 experiments. ****p<0.0001, ***p<0.001, ** p < 0.01, * p < 0.05.

### Co-stimulation of both receptors by 5-HT suppresses 5-HT_2B_-R effect on 5-HT_1A_-R clustering

Since 5-HT_1A_-R and 5-HT_2B_-R respective agonists have differential effects on the membrane clustering of the receptors, we investigated the effect of co-stimulation with their natural ligand, 5-HT (1 µM), on their distribution. Surprisingly, stimulation with 5-HT did not modify 5-HT_2B_-R cluster properties: the cluster density, as well as the cluster size and intensity were not affected by the coexpression of 5-HT_1A_-Rs (**Fig. 8A**). On the contrary, stimulation of 5-HT_1A_-R with 5-HT decreased its surface expression by 39% compared to unstimulated condition (**Fig. 8B**), and close to what was observed after 8-OH-DPAT application (**Fig. 5B**). When coexpressed with 5-HT_2B_-Rs, 5-HT_1A_-R surface expression increased by 24%, and 5-HT stimulation decreased this surface expression by 50%, this effect being fully reversed by the selective 5-HT_2B_-R antagonist, RS (**Fig. 8B**). This result supports the notion that stimulating 5-HT_1A_/5-HT_2B_ heterodimers by 5-HT somehow reverses the 5-HT_2B_-R ability to maintain the 5-HT_1A_-R at the plasma membrane by acting at 5-HT_1A_-Rs (**Fig. 8B**). These data indicate a differential crosstalk according to agonist selectivity, 5-HT vs. 8-OH-DPAT or BW.

**Figure 8:**
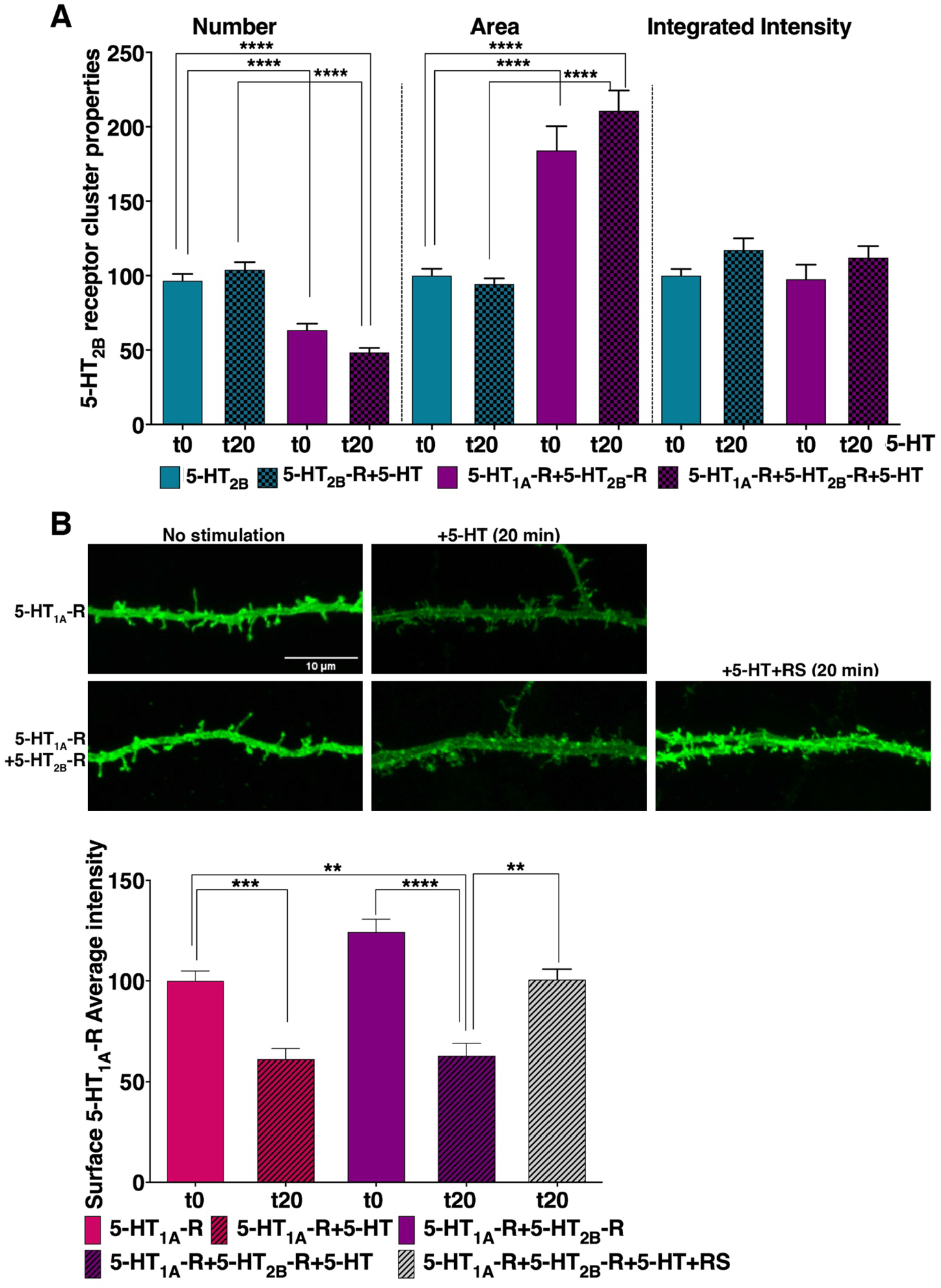
Effect of 5-HT stimulation of 5-HT_1A_-Rs and 5-HT_2B_-Rs expressed separately or in combination on their membrane expression and clustering. (**A**). Quantification of 5-HT_2B_-R membrane clustering following 5-HT stimulation in the presence or not of 5-HT_1A_-Rs. In the presence of 5-HT_1A_-Rs, the number of 5-HT_2B_-R clusters decreased as revealed by Kruskal-Wallis test, P<0.0001, followed by Dunn’s multiple comparisons test; p<0.0001, t0 5-HT_2B_-R vs. t0 5-HT_1A_-R+5-HT_2B_-R and p<0.0001, t0 5-HT_2B_-R vs. t20 5-HT_1A_-R+5-HT_2B_-R, but 5-HT_2B_-R clustering was unaffected upon 5-HT (1 µM) stimulation. In the presence of 5-HT_1A_-Rs, 5-HT_2B_-R cluster area increased, as revealed by Kruskal-Wallis test, P<0.0001, followed by Dunn’s multiple comparisons test, p<0.0001, t0 5-HT_2B_-R vs. t0 5-HT_1A_-R+5-HT_2B_-R and p<0.0001, t0 5-HT_2B_-R vs. t20 5-HT_1A_-R+5-HT_2B_-R, but was unaffected upon 5-HT stimulation; their molecular density (integrated intensity) was not modified by 5-HT stimulation from 0 to 20 min in the absence or presence of 5-HT_1A_-Rs. Data are expressed as percentage of their respective values at t0. (number of cells = 30-39 from 4 experiments). (**B**). Images of 5-HT_1A_-R surface expression acquired by confocal microscopy. Top, 5-HT_1A_-R (green) expression at the dendritic surface in the absence of 5-HT_2B_-Rs and 0 (left), or 20 min (right) after 5-HT stimulation. Bottom, 5-HT_1A_-R expression at the dendritic surface in the presence of 5-HT_2B_-Rs and 0 (left), 20 min (middle) after 5-HT stimulation (1 µM) or 20 min after 5-HT stimulation with the 5-HT_2B_-R antagonist RS (right). Scale bar, 10 µm. Bottom, Quantification of 5-HT_1A_-R surface expression after 5-HT stimulation in the absence or presence of 5-HT_2B_-Rs. The intensity of 5-HT_1A_-Rs at the neuronal surface was decreased by 5-HT stimulation as revealed by Kruskal-Wallis test, P<0.0001, followed by Dunn’s multiple comparisons test, p=0.0003, t0 5-HT_1A_-R vs. t20 5-HT_1A_-R. Similarly, in the presence of 5-HT_2B_-Rs, the 5-HT_1A_-R surface expression was decreased upon 5-HT stimulation, p<0.0001, t0 5-HT_1A_-R+5-HT_2B_-R vs. t20 5-HT_1A_-R+5-HT_2B_-R, but remained unaffected upon 5-HT stimulation in the presence of 5-HT_2B_-R antagonist RS. Number of cells = 27-50, from 4 experiments. Data are expressed as percentage of their respective values at t0. ****p<0.0001, ***p<0.001, ** p < 0.01, * p < 0.05.

### 5-HT_2B_-Rs/5-HT_1A_-Rs control the firing of 5-HT neurons by acting at apamin-sensitive SK channels

The serotonergic tone of the DR neurons is known to be regulated by GIRK channels, which hyperpolarize 5-HT neurons and inhibit cell firing, and by SK channels that are responsible for mAHPs, thereby controlling the burst firing of DR 5-HT neurons. We used patch-clamp recording on *ex-vivo* identified 5-HT neurons to study action potential firing frequency in response to depolarizing current steps in cWT-RCE mice and in cKO^5-HT^-RCE mice. Basal Frequency-Intensity (F-I) curves for cWT and cKO^5-HT^ 5-HT neurons did not differ (**Fig. 9A**), and resting membrane potential and input resistance were similar (**Fig. 9B**). In the presence of 8-OH-DPAT (30 nM), the firing frequency of cKO^5-HT^ 5-HT neurons became significantly higher than that of cWT 5-HT neurons (**Fig. 9C**), while resting membrane potential and input resistance remained similar (**Fig. 9D**). In the presence of 8-OH-DPAT (30 nM) plus Apamin (20 nM), a selective SK2/3 channel blocker, F-I curves for cWT and cKO^5-HT^ 5-HT neurons were again similar (**Fig. 9E**) and higher than unstimulated condition, with resting membrane potential and input resistance remaining similar (**Fig. 9F**). These results indicate that, in the absence of 5-HT_2B_-Rs, 5-HT neuron firing activity is increased upon 5-HT_1A_-R stimulation through the regulation of SK channels, when 5-HT_1A_-R can be internalized by 8-OH-DPAT, and decreased in the presence of 5-HT_2B_-Rs, when 8-OH-DPAT cannot trigger internalization of 5-HT_1A_-Rs, supporting a link between receptor functions and trafficking.

**Figure 9:**
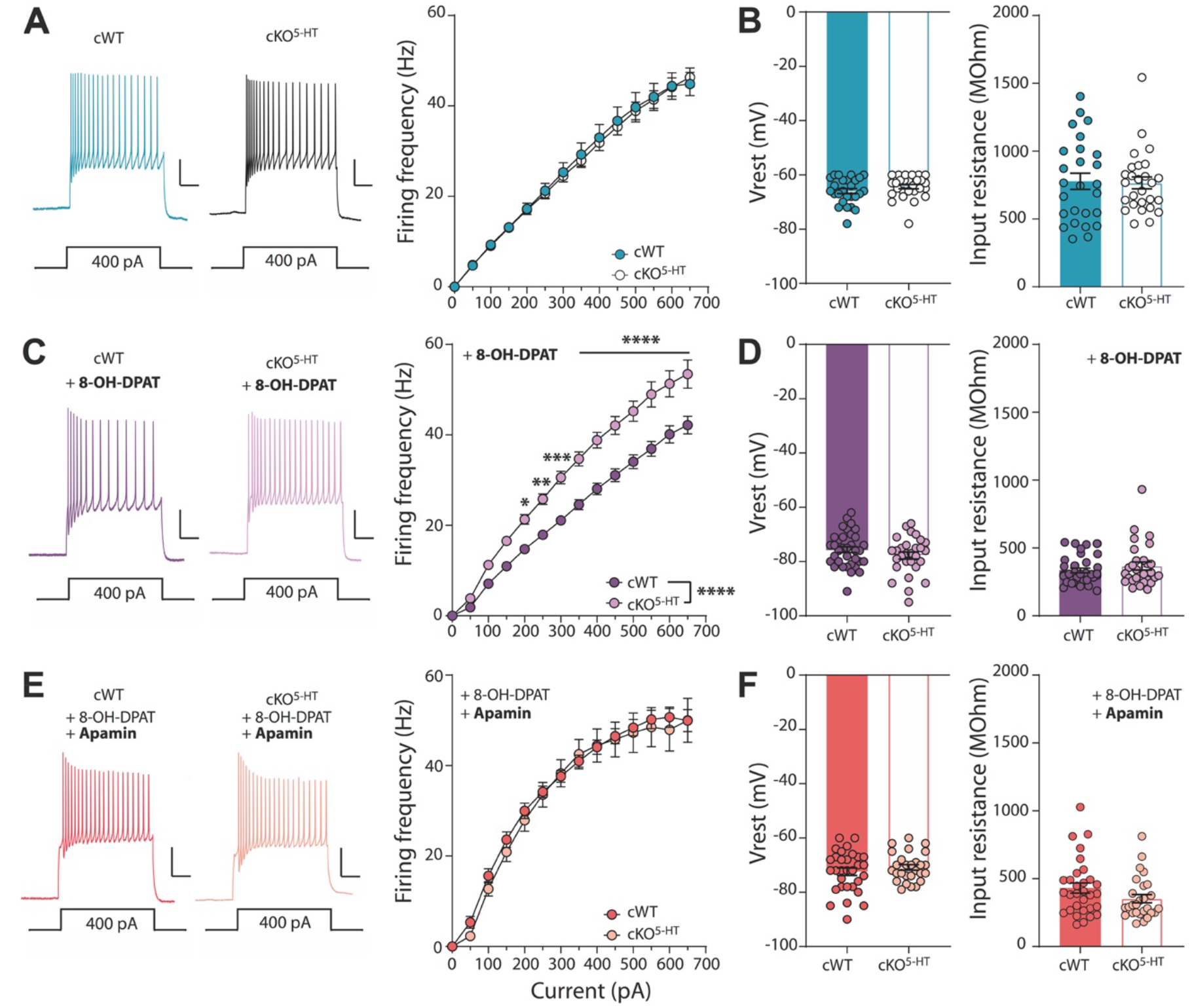
In the absence of 5-HT_2B_-Rs, 5-HT neuron firing activity is increased upon 5-HT_1A_-R stimulation through regulation of SK channels. (**A**). Left, sample spike trains evoked by a 400pA current injection in slices from cWT-RCE and cKO^5-HT^-RCE mice; scale: 25pA/100ms. Right, F-I curve (mean ± SEM) for cWT (n = 26 cells from 7 mice) and cKO^5-HT^ 5-HT neurons (n = 27 cells from 6 mice); two-way RM ANOVA, genotype effect, not significant *p* = 0.8715. **(B**). Resting membrane potential (Vrest) and input resistance in cWT and cKO^5-HT^ 5-HT neurons. Bars represent the mean ± SEM; circles show individual recordings. (**C**). Left, same as in (**A**) in presence of 8-OH-DPAT (30 nM), a 5-HT_1A_-R selective agonist. Right, F-I curve (mean ± SEM) for cWT (n = 35 cells from 5 mice) and cKO^5-HT^ 5-HT neurons (n = 29 cells from 4 mice) in the presence of 8-OH-DPAT (30 nM); two-way RM ANOVA revealed a significant increase in firing rate of cKO^5-HT^, genotype effect p<0.0001; Sidak’s *post hoc* test* p=0.0274, ** p=0.0028, *** p=0.0001, **** p<0.0001. (**D**) Same as in (B) in the presence of 8-OH-DPAT (30 nM). (**E**) Left, same as in (**A**) and (**C**) in the presence of 8-OH-DPAT (30nM) and Apamin (20 nM), a selective SK channel blocker. Right, F-I curve (mean ± SEM) for cWT (n = 31 cells from 4 mice) and cKO^5-HT^ 5-HT neurons (n = 27 cells from 3 mice) in the presence of 8-OH-DPAT (30 nM) and Apamin (20 nM); two-way RM ANOVA, no genotype effect *p* = 0.6979. (**F**) Same as in (**B**) and (**D**) in the presence of 8-OH-DPAT (30 nM) and Apamin (20 nM).

## DISCUSSION

Combined with our previous investigations about 5-HT_2B_-Rs functions in 5-HT neurons (Belmer et al., 2018) and the extensive studies on 5-HT_1A_-Rs as autoreceptors of these neurons, the present work demonstrates 5-HT_1A_-R/5-HT_2B_-R interactions are implicated in the regulation of serotonergic tone, which is a prime importance for mood disorders. We found that 5-HT_1A_-Rs and 5-HT_2B_-Rs are colocalized in 5-HT neurons, and interact together to form dimers. This interaction modifies their cell-surface expression and distribution, which can be regulated upon response to specific ligand.

Different techniques allowed us to validate the reciprocal effects on surface 5-HT_1A_-R and 5-HT_2B_-R. Confocal microscopy was used to analyze and compare in various conditions the number of clusters, their surface and the density of receptors per cluster. With a resolution up to ∼20 nm, STORM allowed an accurate analysis of receptor spatial distributions (MacDonald et al., 2015). This technique validated that 5-HT_2B_-R stimulation by a selective 5-HT_2B_-R agonist, BW, triggered the formation of larger and denser clusters of 5-HT_2B_-Rs at the surface of hippocampal neurons (**Fig. 10A**). 5-HT_1A_-R coexpression significantly decreased the 5-HT_2B_-R cluster number, increased their area, and potentiated these effects upon 5-HT_2B_-R stimulation by BW (**Fig. 10B**). Coexpression with 5-HT_2B_-R increases 5-HT_1A_-R expression and maintains 5-HT_1A_-Rs at the neuronal surface despite its own stimulation by 8-OH-DPAT (**Fig. 10B**). Selective stimulation by 8-OH-DPAT of 5-HT_1A_-Rs expressed alone decreases their surface expression (**Fig. 10A**). It has been shown that the 5-HT_1A_ autoreceptor desensitizes and is internalized upon stimulation (Bouaziz et al., 2014) and especially after chronic exposure to high extracellular 5-HT as upon exposure to SSRIs (Soiza-Reilly et al., 2015), although the detailed mechanisms involved remained unclear. Phosphorylation sites were identified in human 5-HT_1A_-Rs. Activation of PKA leads to phosphorylation and desensitization of the 5-HT_1A_-R with a stoichiometry of one phosphate per receptor (Raymond et al., 1999), and stimulation of PKA increases the PKC-induced phosphorylation and desensitization. Using BRET (Stroth et al., 2015), 5-HT_1A_-Rs was shown to recruit β-arrestin2 that co-internalized with 5-HT_1A_-Rs upon agonist stimulation (Albert and Vahid-Ansari, 2019). An uncoupling of 5-HT_1A_-Rs from G-proteins and from the inhibition of adenylyl cyclase was also associated to the actions of GRKs (Nebigil et al., 1995) since coexpression of GRK2 and β-arrestin2 triggered 5-HT_1A_-R internalization via clathrin-coated pits (Heusler et al., 2008), and signaling to ERK1/2 (Della Rocca et al., 1999). Concerning 5-HT_2B_-Rs, we showed that 5-HT_2B_-R internalization was dependent on clathrin and β-arrestin-2, but independent of Caveolin-1 (Cav1) and GRK2,3 when stimulated by 5-HT (Janoshazi et al., 2007), whereas stimulation by BW internalized 5-HT_2B_-Rs more than twice faster than 5-HT and relied on GRK2,3, clathrin, and β-arrestin2, independently of Cav1. Similar transduction mechanisms likely take place in 5-HT neurons but remain to be documented.

**Figure 10:**
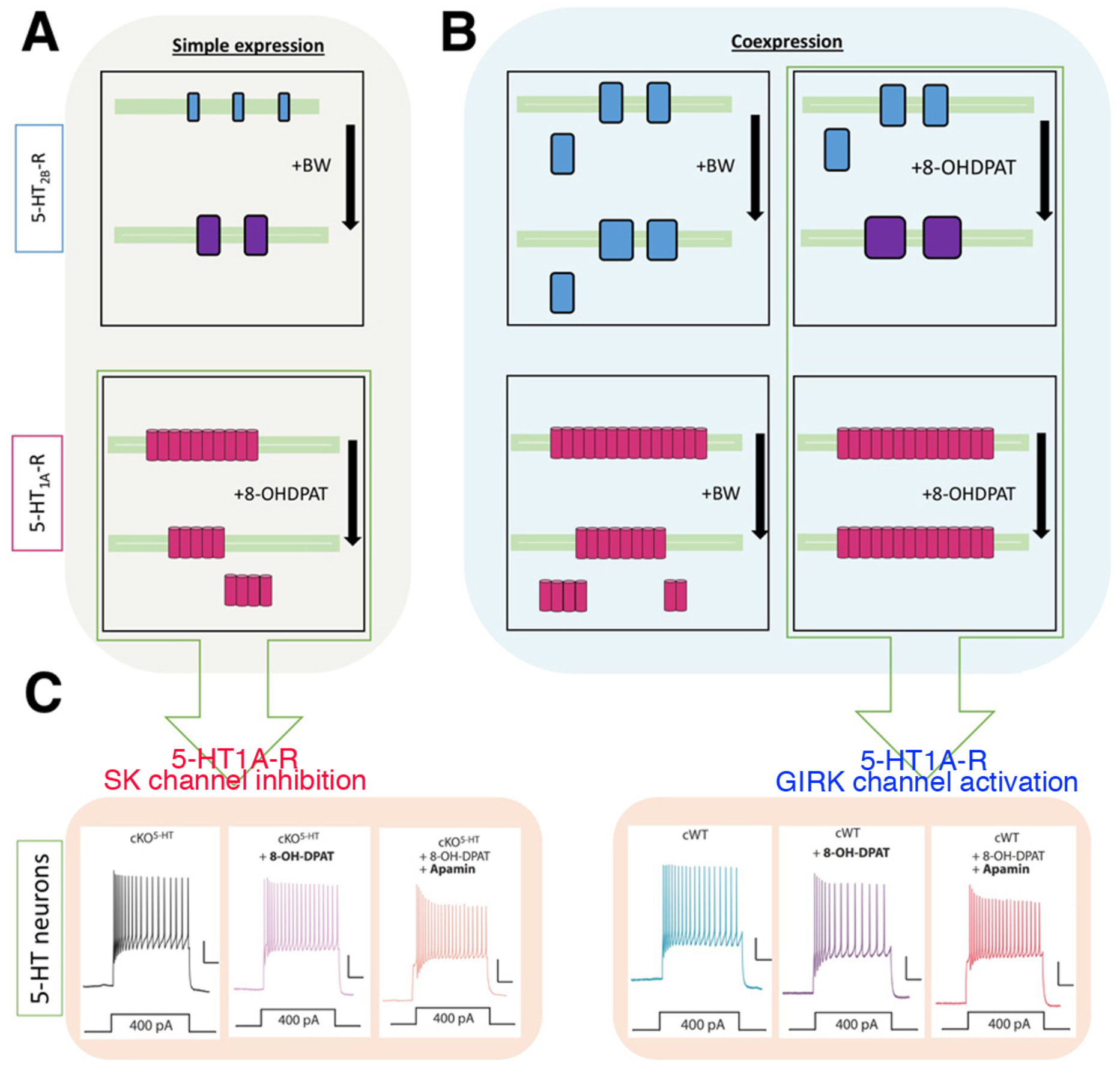
Effect of 5-HT_1A_-R and 5-HT_2B_-R co-expression on their membrane expression, clustering, and functional impact. (**A**). Separate expression of 5-HT_1A_-Rs or 5-HT_2B_-Rs. Top, BW stimulation increases 5-HT_2B_-R clustering. Bottom, 8-OH-DPAT stimulation decreases surface expression of 5-HT_1A_-Rs probably via endocytosis. (**B**). Co-expression of 5-HT_2B_-Rs and 5-HT_1A_-Rs and their stimulation affects the membrane expression of both receptors. Top, 5-HT_1A_-R expression increases 5-HT_2B_-R clustering. The 5-HT_2B_-R stimulation by BW does not reverse the clustering effect of 5-HT_1A_-Rs on 5-HT_2B_-Rs. Stimulation by 8-OH-DPAT of 5-HT_1A_-Rs further increases 5-HT_2B_-R clustering. Bottom, Coexpression of 5-HT_2B_-Rs increases 5-HT_1A_-R surface expression. No further change in surface expression of 5-HT_1A_-Rs is detected upon its stimulation with 8-OH-DPAT, but 5-HT_2B_-R stimulation by BW decreases 5-HT_1A_-R surface expression. (**C**). The 5-HT_2B_-R/5-HT_1A_-R co-expression/interaction regulates the excitability of 5-HT neurons. In the absence of 5-HT_2B_-Rs (Separated expression, i.e. cKO^5-HT^, left), 5-HT neuron firing activity is increased upon 5-HT_1A_-R stimulation by 8-OH-DPAT, independently of SK channels blockade by apamin. By contrast, upon co-expression of 5-HT_2B_-R s(Co-expression, i.e. cWT, right), 5-HT neuron firing activity is decreased by 8-OH-DPAT unless SK channels are blocked by apamin.

Oligomerization is an important process for GPCRs and can be essential for their trafficking to the plasma membrane, agonist binding, G-protein activation and signaling (Herrick-Davis, 2013; Faron-Górecka et al., 2019). 5-HT-Rs are able to form homo or heterodimers with other 5-HT-Rs or non-5-HT-Rs (Maroteaux et al., 2019). 5-HT_1A_-R/5-HT_7_-R heterodimerization decreases Gαi activation and inhibits GIRK activation, which are both mediated by 5-HT_1A_-Rs without affecting the 5-HT_7_-R signaling (Renner et al., 2012). 5-HT_1A_-Rs and 5-HT_2A_-Rs (close relative to 5-HT_2B_-Rs) can form oligomers (Borroto-Escuela et al., 2017), and in pyramidal neurons of the prefrontal cortex, coexpression of these two receptors triggers opposite and mixed effects of excitation/inhibition (Avesar and Gulledge, 2012; Ju et al., 2020). In cells coexpressing 5-HT_1B_-Rs (close relative to 5-HT_1A_-Rs) and 5-HT_2B_-Rs, and in the absence of direct interaction, their co-stimulation by 5-HT makes 5-HT_2B_-R to internalize via its classic clathrin pathways but also via 5-HT_1B_-triggered GRK2,3 activation and Caveolin1-dependent pathways (Janoshazi et al., 2007). Here, we show that coexpression of 5-HT_1A_-Rs and 5-HT_2B_-Rs, even if their interaction has no apparent impact on their classical second messenger signalization, has an effect on the expression levels of both receptors at the plasma membrane (**Fig. 10B**). The specific interacting sequence in the heteromer between the 5-HT_2B_-R and the 5-HT_1A_-R has now to be identified.

We previously reported that *in-vivo* injection of BW attenuated 8-OH-DPAT-induced hypothermia and firing inhibition of 5-HT neurons, both of which being attributed to the action of 5-HT_1A_ autoreceptors on 5-HT neurons. These data supported that 5-HT_2B_-Rs were able to counteract 5-HT_1A_-R negative effects (Belmer et al., 2018). In the present study, we reveal that both receptors are expressed in the same neuronal somatodendritic compartment of raphe 5-HT neurons and that they physically and functionally interact. One important function of 5-HT_1A_-Rs is the Gβγ-dependent activation of GIRK channels, which leads to hyperpolarization and reduction of 5-HT neuron firing (Lüscher et al., 1997; Llamosas et al., 2015; Montalbano et al., 2015). The 5-HT_1A_-R/5-HT_2B_-R interaction could thus modify Gβγ-dependent outcomes. It has been shown that activation of GIRK channels relies on promiscuous presence of PIP2 (Niu et al., 2020). However, the activation of 5-HT_2B_-R promotes IP production through activation of PLC (Maroteaux, 2021). Since these two receptors interact within the same membrane microdomain, activation of 5-HT_2B_-R could prevent 5-HT_1A_-R signal transduction cascade to GIRKs. This hypothesis could explain the previously mentioned *in-vivo* observations about hypothermia and firing (Belmer et al., 2018). Although the mechanism underlying the modulation of 5-HT neuron activity by 5-HT_2B_-Rs remains to be clarified, the present study provides initial insight into the regulation of 5-HT_1A_-R surface expression by 5-HT_2B_-Rs through their interactions as dimers specifically in 5-HT neurons, thereby modulating serotonergic neuron activity.

Since coexpression of 5-HT_1A_-Rs with 5-HT_2B_-Rs mimics clustering effects of 5-HT_2B_-R stimulation, and is enhanced by selective 5-HT_1A_-R stimulation with 8-OH-DPAT, we conclude that 5-HT_1A_-R expression without or with stimulation by 8-OH-DPAT can control 5-HT_2B_-R clustering. Reciprocally, while homologous 5-HT_1A_-R agonist stimulation by 8-OH-DPAT decreases 5-HT_1A_-R surface expression, coexpression with 5-HT_2B_-R increases 5-HT_1A_-R expression and maintains 5-HT_1A_-Rs at the neuronal surface despite its own stimulation by 8-OH-DPAT. Therefore, the dynamic regulation of 5-HT_1A_-R surface expression not only depends on the ligand but also on cell types and their receptor expression. In addition, we show that 5-HT, the physiological ligand of 5-HT_1A_-Rs and 5-HT_2B_-Rs, has no significant impact on the distribution of 5-HT_2B_-Rs coexpressed with 5-HT_1A_-Rs. Interestingly, we found that 5-HT stimulation abolishes 5-HT_2B_-R positive effect on 5-HT_1A_-R surface expression, and decreases 5-HT_1A_-R surface expression level. This decrease was entirely 5-HT_2B_-R-dependent since it can be restored by applying the 5-HT_2B_-R antagonist RS, coherent with the observation that BW reduces 5-HT_1A_-R surface expression only in the presence of 5-HT_2B_-Rs. Of note, PKC, which can be activated by 5-HT_2B_-Rs was shown to induce a rapid phosphorylation of 5-HT_1A_-Rs at a stoichiometry of two phosphates per receptor (Raymond et al., 1999), and to preferentially uncouple Gβγ over Gαi signaling (Wurch et al., 2003; Kushwaha et al., 2006). Together, these results demonstrate that modulation of 5-HT_1A_-R surface expression can be triggered not only by its own stimulation, but also through 5-HT_2B_-R activation, confirming their functional crosstalk and explaining, at least in part, the *in vivo* interaction observed between these two receptors. The difference between effects of 5-HT_2B_-R stimulation by 5-HT and BW may be explain by its biased coupling to GRK2,3 that we previously reported (Janoshazi et al., 2007). We finally confirmed the lower efficacy of 5-HT, which had no obvious clustering or internalization effect on 5-HT_2B_-Rs in the presence of 5-HT_1A_-Rs.

We previously showed by cell-attached electrophysiological recording that activation of 5-HT_2B_-Rs by BW did increase 5-HT neurons firing activity (Belmer et al., 2018). Here, using current-clamp in *ex-vivo* identified 5-HT neurons, we observed that the increase in firing frequency of action potentials upon raising current steps was similar in cWT mice as in cKO^5-HT^ mice. Surprisingly, stimulation by 8-OH-DPAT increased significantly more firing frequency of 5-HT neurons in the absence of 5-HT_2B_-Rs in cKO^5-HT^ mice than in 5-HT neurons from cWT mice, i.e. in the presence of 5-HT_2B_-Rs and 5-HT_1A_-Rs. Only the inhibition of SK channels with apamin allowed 5-HT neurons from cWT mice to fire as 5-HT neurons from cKO^5-HT^ mice. This work, therefore, supports the possibility that 5-HT_2B_-Rs somehow compete with 5-HT_1A_-Rs to inhibit SK channels. Previously, using current-clamp recordings we showed that a simple overexpression of 5-HT_2B_-Rs in 5-HT neuron triggered a significant increase in the basal (non-stimulated) firing frequency compared to controls (Belmer et al., 2018). Apamin was shown to increase the burst firing of DR 5-HT neurons leading to increased 5-HT release (Crespi, 2009) in a similar way as the SSRI fluoxetine (Crespi, 2010) or as the 5-HT_2B_-R agonist BW (Diaz et al., 2012; Belmer et al., 2018). Furthermore, mice exposed to chronic social isolation develop depressive-like state that can be reversed by SSRIs only in the presence of 5-HT_2B_-Rs (Diaz et al., 2016), or by apamin, which regulates 5-HT firing pattern (Sargin et al., 2016). Therefore, interaction between 5-HT_2B_-R/5-HT_1A_-R and their stoichiometry are required to control proper firing of 5-HT neurons by regulating mAHP that conditioned SSRIs efficacy.

These functional results support our findings on 5-HT_1A_-R/5-HT_2B_-R interactions. The coexpression of 5-HT_1A_-Rs with 5-HT_2B_-Rs increases 5-HT_1A_-R membrane expression by blocking its internalization by 8-OH-DPAT and also prevents 5-HT_1A_-R coupling to inhibition of SK channels but favors its activation of GIRK channels. Here, only the exposure to 8-OH-DPAT together with apamin is able to trigger in cWT neurons a firing response similar to that of cKO^5-HT^ neurons, supporting a loss of ability of 5-HT_1A_-Rs to couple to SK channel inhibition in the presence of 5-HT_2B_-Rs. Our previous report that either 5-HT_2B_-R overexpression or its stimulation by BW was able to increase 5-HT neuron firing (Belmer et al., 2018) together with the present observation that coexpression with 5-HT_1A_-Rs increases the size of 5-HT_2B_-R clusters potentiated by 8-OH-DPAT, supports 5-HT_2B_-R ability to couple to SK channel inhibition in this condition. By contrast, in the absence of 5-HT_2B_-Rs, 8-OH-DPAT triggers internalization of 5-HT_1A_-Rs, and favors its coupling to SK inhibition (**Fig. 10C**) as previously reported (Grunnet et al., 2004).

Together, these data support that the ratio between 5-HT_1A_-R and 5-HT_2B_-R expression tunes the firing pattern of 5-HT neurons through SK and GIRK channels regulation and thus conditions the SSRI efficacy. It remains to be determined if 5-HT_2B_-R stimulation controls directly (e.g. by PKC activation and thus by phosphorylation) SK channels, or if this regulation is relying exclusively on 5-HT_1A_-R/5-HT_2B_-R dimeric interactions and competition to common effectors.

## MATERIALS AND METHODS

### Generation and use of mice

Floxed mice, *Htr2b*^*tm2Lum*^*/Htr2b*^*tm2Lum*^, (IMSR Cat# EM:05939, RRID: IMSR_EM:05939) (cWT) were generated on a mixed B6;129S2 background and backcrossed >10 times onto the 129S2 strain. *Htr2b*^*lox/lox*^ mice were inactivated for *Htr2b* in 5-HT neurons by crossing with *129S2*.*Cg-Tg(Fev-cre)*^*1Esd*^*/0* (IMSR Cat# JAX:012712, RRID:IMSR_JAX:012712) (ePet1-Cre BAC transgenic mice or *Pet1-Cre*^*+/0*^) (Scott et al., 2005) generating *129S2*.*Cg-Pet1-Cre*^*+/0*^; *Htr2b*^*lox*^*/Htr2b*^*lox*^ conditional knockout mice (cKO^5-HT^) and littermate controls (cWT) (Belmer et al., 2018). For electrophysiology, these strains of mice were further crossed with *Gt(ROSA)26Sor*^*tm1(CAG-EGFP)Fsh*^*/Gt(ROSA)26Sor*^*+*^ (MGI Cat# 4420760, RRID:MGI:4420760) (Rosa26;CAG-loxP-STOP-loxP-EGFP or RCE) that after crossing with Pet1-Cre^+/0^ generated (Pet1-GFP) that express GFP in Pet1-positive 5-HT neurons only after Cre recombination cWT-RCE, and in the presence of floxed allele generated conditional knockout mice (cKO^5-HT^-RCE) that express GFP in Pet1-positive 5-HT neurons.

All mice were bred at our animal facility. Food and water were available *ad libitum*. The temperature was maintained at 21±1°C, under 12/12h light/dark. Mice were moved to the experimental room in their home cage at least 5 days prior to testing to allow for habituation to the environment. All experiments involving mice were approved by the local ethical committee, Ethical Committee for Animal Experiments of the Sorbonne University, Charles Darwin C2EA - 05 (authorizations N°APAFIS#28228, and APAFIS#28229). All efforts were made to minimize animal suffering. Results are described in accordance with the ARRIVE guidelines for reports in animal research (Percie du Sert et al., 2020).

### Experimental design and statistical analysis

Statistical analyses were performed using GraphPad Prism 7 software. Starting from our experience in previous publications (Belmer et al., 2018; Benhadda et al., 2021), we calculated the standard deviation of each group of animals for each test as well as the differences between the means of these groups. By setting the type I and II error risks at 5 and 20%, respectively, the size of the groups were determined by calculating the statistical power using the G*Power software version 3.1.9.6. To ensure reproducibility, when relevant, experiments were performed at least three times independently. To avoid litter bias in the mouse experiments, experimental groups are composed of animals from different litters randomly distributed using Graphpad software. All analyses were conducted with blinding to the experimental condition. Putative outliers were determined by the ROUT method. Data are presented as bars and the line at the mean ± S.E.M. (standard error of the mean). Comparisons between two groups following a normal distribution were analyzed using two-tailed unpaired t-test with or without Welch’s correction. Normality was assessed using the D’Agostino & Pearson omnibus normality test. All data normally distributed for more than two groups were examined by either ordinary or repeated measure two-way ANOVA followed by Bonferroni’s *post hoc* comparisons. P<0.05 was predetermined as the threshold for statistical significance (See supplementary data for full statistical analysis).

### Viral constructs and stereotaxic injection

To express HA-tagged 5-HT_2B_-R and Flag-tagged 5-HT_1A_-R specifically in 5-HT neurons, we use a Double floxed Inverse Orientation (DIO) adeno-associated virus (AAV) construct that allows Cre-mediated expression of the transgene (pAAV-EF1A-DIO-WPRE-pA vector; RRID Addgene_39320) (Belmer et al., 2018). The viruses packaged into AAV2.9 serotype with titers of 10^12^–10^13^ viral particles / ml were obtained (UNC Vector Core, Chapel Hill USA). AAVs were injected in the B7 raphe nuclei of *Pet1-Cre*^*+/0*^ mice following described procedure (Muzerelle et al., 2016). Respective expression was assessed by immunostaining using a mouse anti-HA antibody (1:500; Cell Signaling#2367, RRID AB_10691311 and a rabbit anti-Flag (1:500; Cell Signaling#14793 RRID AB_2572291 followed by Cy5-conjugated donkey anti-mouse (1.9 µg/ml; Jackson ImmunoResearch#715-175-150, RRID AB_2340819) and the Cy3-conjugated goat anti-rabbit antibody (1.9 µg/ml; Jackson ImmunoResearch#111-165-003, RRID AB_2338000).

### Culture of COS-7 cells

COS-7 cells were cultured as monolayers in Dulbecco’s modified Eagle’s medium (DMEM) (Gibco, Invitrogen) supplemented with 10% fetal calf serum (Biowest) and 1% penicillin/streptomycin (Sigma), in 9-cm dishes (Falcon). Cells were incubated at 37 °C in a 5% CO_s_ atmosphere. Cells 70% confluent in 6-well plates were used for inositol phosphate accumulation and surface biotinylation, in 24-well plates for surface binding, in 9-cm dishes for total binding and co-immunoprecipitation/Western-blot and in 96-well plates for BRET experiments. They were transfected with pCAGG vectors using the Genejuice transfection reagent (Merck Millipore) in complete DMEM, according to the manufacturer’s instructions. Twenty-four hours before the experimental test, cells were incubated in a serum-free medium.

### Co-immunoprecipitation

Co-immunoprecipitation and Western-blotting were performed as previously described (Moutkine et al.). COS-7 cells transfected with a total of 10 µg of DNA per 9-cm dishes with a ratio of 7:3 (HA-5-HT_2B_ / empty vector; empty vector / myc-5-HT_1A_; HA-5-HT_2B_ / myc-5-HT_1A_) and maintained in culture for three days before harvesting. Cells were centrifuged and suspended in CHAPS lysis buffer (50 mM Tris-HCl, pH 7.4, 0.05 mM EDTA, 10 mM CHAPS, and protease inhibitor cocktail, pH 7.4) and sonicated three times for 30 s. Cells were next solubilized for 5 h at 4°C under gentle agitation. Lysates were centrifuged (20,000 x g) in order to pellet non-solubilized membranes. Protein concentrations in supernatant were measured using the PierceTM Coomassie Protein Assay Kit. Lysates were co-imunoprecipitated with anti-HA beads (HA-Tag C29F4 Rabbit mAb Sepharose^®^ Bead Conjugate Cell Signaling #3956 RRID:AB_10695091) overnight at 4 °C under gentle agitation. Total lysate and immunoprecipitated proteins were separated by SDS/PAGE onto 10% polyacrylamide gels and transferred electrophoretically to nitrocellulose membranes. Inputs represent 5% of the total protein amount used for immunoprecipitations. Blots were probed with rabbit anti-HA (1:1,000; Cell Signaling#3724, RRID AB_1549585) or mouse anti-myc (1:1,000; Cell Signaling#2276, RRID AB_331783). Antibodies (1:10,000; goat anti-mouse-IR800, Advansta# R-05060-250 and goat anti-rabbit-IR700, Advansta# R-05054-250) were used as secondary antibodies. Immunoreactive bands were detected using the Odyssey software. At least three independent experiments were performed.

### Bioluminescence resonance energy transfer (BRET)

BRET assays were performed according to published methods (Moutkine et al., 2017). Briefly, in 6-well plates, COS-7 cells were transfected with 30-100 ng of plasmid DNA coding for the BRET donor (5-HT_2B_-R-Rluc) (Moutkine et al., 2017) and an increasing amount of BRET acceptor plasmids (5-HT_1A_-R-YFP; from 100 to 4000 ng/well). Twenty-four hours later, the cells were trypsinized (trypsin 0.05% EDTA; Invitrogen) and plated in 96-well plates (50,000 cells/well). The day after, the luciferase substrate, coelenterazine-h, was added in each well at a 5 µM final concentration. Using a Mithras LB940 plate reader, luminescence and fluorescence were then measured simultaneously at 485 and 530 nm, respectively. The BRET ratios were calculated as ((emission at 530 nm/emission at 485 nm)-(background at 530 nm/background at 485 nm)) and were plotted as a function of ((YFP-YFP0)/YFP0)/(Rluc/Rluc0)). The background corresponds to signals in cells expressing the Rluc fusion protein alone under the same experimental conditions. YFP is the fluorescence signal at 530 nm after excitation at 485 nm, and Rluc is the signal at 485 nm after addition of coelenterazine-h. YFP0 and Rluc0 correspond to the values in cells expressing the Rluc fusion protein alone.

### Binding experiments

#### Membrane radioligand binding assay

Membrane binding assays were performed on transfected cells plated in 9-cm dishes (Belmer et al., 2018). Cells were first washed with PBS, scraped into 10 ml of PBS on ice, and then centrifuged for 5 min at 1000 g. Cell pellets were dissociated and lysed in 2 ml of binding buffer (50 mM Tris-HCl, 10 mM MgCl_2_, 0.1 mM EDTA, pH 7.4) and centrifuged for 30 min at 10,000 g. Membrane preparations were then resuspended in binding buffer to obtain a final concentration of 0.2–0.4 mg of protein/well. Aliquots of membrane suspension (200 µl/well) were incubated with 25 µl/well of ^3^H-mesulergine or ^3^H-8-OH-DPAT at a final concentration between 1/2 to 1/10 Kd for each 5-HT-R, diluted in binding buffer and 25 µl/well of increasing concentrations of heterologous compound. 5-HT_2B_-R agonist BW-723C86 (BW) or antagonist RS-127445 (RS), 5-HT_1A_-R agonist 8-OH-DPAT or antagonist NAN-190 (NAN) were from Tocris (UK). Competition was performed at concentration between 10^−11^ to 10^−5^ M, diluted in binding buffer in 96-well plates for 60 min at room temperature. Membranes were harvested by rapid filtration onto Whatman GF/B glass fiber filters (Brandell) pre-soaked with cold saline solution and washed three times. Filters were placed in 6-ml scintillation vials and counted. Data in disintegrations/min were converted to femtomoles and normalized to protein content (ranging from 0.1 to 1 mg/well). At least three independent experiments were performed in duplicate.

#### Non-permeabilized whole cell radioligand binding assay

Cells expressing 5-HT_2B_-Rs and/or 5-HT_1A_-Rs were plated in 24-well clusters. Twenty-four hours before the experiment, the cells were incubated in serum-free medium overnight. The next day, the medium was replaced by 400 µl/well of Krebs-Ringer/Hepes buffer (130 mM NaCl, 1.3 mM KCl, 2.2 mM CaCl_2_, 1.2 mM NaH_2_PO_4_, 1.2 mM MgSO_4_, 10 mM Hepes, 10 Mm glucose, pH 7.4). Then, 50 µl of [^3^H]-mesulergine were diluted in Krebs-Ringer/Hepes buffer at a final concentration 1/2 Kd value of 5-HT_2B_-R. The radioligand was competed with 50 µl of increasing concentrations of non-radioactive BW, also diluted in Krebs-Ringer/Hepes buffer. Cells were then incubated for 30 min at room temperature and then washed twice on ice with cold PBS. Washed cells were solubilized by the addition of 500 µl of SDS 1%. The next day, 3 ml of scintillation mixture were added to the samples, and the radioactivity was counted. Data in disintegrations/ min were converted to femtomoles and normalized to protein content (0.2–0.4 mg of protein/well). At least three independent experiments were performed in duplicate.

### Second messenger measurements

COS-7 cells were transfected with 3 µg of DNA (1:1 ratio for co-transfection) in 6-well plates using Genejuice transfecting reagent in complete medium. Twenty-four hours later the cells were trypsinized (trypsin 0.05% EDTA; Invitrogen) and plated in 96-well plates (30,000 cells/well). The next day, the complete medium was replaced by a serum-free medium.

#### HTRF IP accumulation

The day of the experiment, media were replaced by stimulation buffer with LiCl to prevent IP1 degradation (NaCl, 146 mM, KCl, 4.2 mM, MgCl_2_, 0.5 mM, CaCl_2_, 1 mM, Hepes, 10 mM, glucose, 5.5 mM, LiCl, 50 mM, pH7.4). Cells were stimulated during 2 h at 37°C with different concentrations of BW (10^−11^ to 10^−6^ M in stimulation buffer). Stimulation solution was replaced by a lysis buffer (IP one HTRF Kit, Cisbio, France) during 1 h. Lysates were distributed to 384-well plates, and IP was labeled using HTRF reagents. The assay is based on a competitive format involving a specific antibody labeled with terbium cryptate (donor) and IP coupled to d2 (acceptor). After a 1-h incubation with HTRF reagent, the plate was read using Mithras LB940 plate reader according to the manufacturer’s instructions. Modelization and EC50 calculation were done using GraphPad Prism 7 software. At least three independent experiments were performed in duplicate.

### Surface biotinylation

Surface biotinylation was performed on transfected COS-7 cells. Briefly, cells were initially washed twice in cold PBS and subsequently incubated in PBS containing 1 mg/ml Sulfo-NHS-SS-Biotin for 30 min at 4°C to allow for labelling of all surface membrane proteins. Reaction was stopped by applying a quench solution (100 mM Tris pH 8 in PBS) to remove excess biotin. Biotinylated cells were then homogenized in RIPA buffer containing 25 mM Tris (pH 7.6), 150 mM NaCl, 1% TritonTM X-100, 0.5% sodium deoxycholate, 0.1% SDS, 1mM NaF and a cocktail of protease inhibitors (Roche). The lysate was centrifuged at 21,000 g to remove nuclei and cellular debris. A small amount of the lysate was removed and constituted the “input” or total lysate. Then, 100 µl of StreptaAvidin beads (Thermo Scientific Inc) were added to 40 µg of protein lysate and placed on a rotator at 4°C overnight. Samples were then washed three times in RIPA buffer and a fourth times in RIPA buffer without detergents (25 mM Tris pH 7.6, 150 nM NaCl, 0.5% TritonTM X-100; protease inhibitor); beads were pulled-down after each wash by 1 min centrifugation. Bound proteins were eluted in SDS reducing buffer and heated at 70°C for 5 min.

### Western blotting

For co-immunoprecipitation and surface biotinylation experiments, total lysate and immunoprecipitated proteins were separated by SDS/PAGE onto 10% acrylamide gels and transferred electrophoretically to nitrocellulose membranes. Blots were probed with rabbit anti-HA (1:1,000; Cell Signaling#3724, RRID AB_1549585) or mouse anti-myc (1:1,000; Cell Signaling#2276, RRID AB_331783). Secondary antibodies (1:10,000; goat anti-mouse-IR800, Advansta# R-05060-250 and goat anti-rabbit-IR700, Advansta# R-05054-250) were used as secondary antibodies. Immunoreactive bands were detected using the Odyssey software. At least three independent experiments were performed.

### Hippocampal neuronal culture and transfection

Hippocampal neurons were prepared from embryonic day 19 Sprague-Dawley rat pups as previously described (Benhadda et al., 2021). Dissected tissues were trypsinized (0.25% v/v) and mechanically dissociated in HBSS (CaCl_2_ 1.2 mM, MgCl_2_ 0.5 mM, MgSO_4_ 0.4 mM, KCl 5 mM, KH_2_PO_4_ 0.44 mM, NaHCO_3_ 4.1 mM, NaCl 138 mM, Na_2_HPO4 0.34 mM, D-Glucose 5.5 mM) containing 10 mM Hepes (Invitrogen). Dissociated cells were plated on glass coverslips (Assistent) precoated with 55 µg/ml poly-D,L-ornithine (Sigma-Aldrich) in plating medium composed of MEM supplemented with horse serum (10% v/v; Invitrogen), L-glutamine (2 mM), and Na pyruvate (1 mM, Invitrogen) at a density of 3.4 × 10^4^ cells/cm^2^ and maintained in humidified atmosphere containing 5% CO_2_ at 37°C. After attachment for 2-3 h, cells were incubated in maintenance medium that consists of Neurobasal medium supplemented with B27, L-glutamine (2 mM), and antibiotics (Invitrogen). Each week, one-third of the culture medium volume was renewed. Neuronal transfection with plasmids encoding HA-5-HT_2B_-R, Myc-5-HT_1A_-R and eGFP were performed at 13–14 DIV using Transfectin (Bio-Rad), according to the instructions of the manufacturer (DNA/lipofectant ratio of 1:3), with 1.5 µg of plasmid DNA per 20-mm well. The following ratio of plasmid DNA was used in co-transfection experiments: 1:0.3:0.2 µg for HA-5-HT_2B_-R / Myc-5-HT_1A_-R / eGFP. Experiments were performed 7–10 days after transfection.

### Immunostaining

The total (membrane plus intracellular) pools of HA-5-HT_2B_-Rs and myc-5-HT_1A_-Rs were revealed with immunocytochemistry in fixed and permeabilized cells, whereas the membrane pool was done in non-permeabilized cells. To label the total pool of receptors, cells were fixed for 10 min at room temperature in paraformaldehyde (PFA; 4% w/v; Sigma) and sucrose (30% w/v; Sigma) solution in PBS. Cells were then washed in PBS, and permeabilized for 4 min with Triton X-100 (0.25% v/v) in PBS. After these washes, non-specific staining was blocked for 30 min with goat serum (20% v/v; Invitrogen) in PBS. Neurons were then incubated for 1 h with either the rabbit anti-HA antibody (1:500; Cell Signaling #3724, RRID AB_1549585) or the mouse anti-myc antibody (1:500; Cell Signaling #2276, RRID AB_331783) in PBS supplemented with Goat Serum (GS) (3% v/v). Cells were then washed three times and incubated for 45 min with Cy5-conjugated donkey anti-mouse or Alexa 488 donkey anti-mouse antibodies (1.9 µg/ml; Jackson ImmunoResearch # 715-175-150, RRID AB_2340819 or #715-175-150, RRID AB_2336933) or the Cy3-conjugated goat anti-rabbit antibody (1.9 µg/ml; Jackson ImmunoResearch # 111-165-003, RRID AB_2338000) in PBS-BSA-goat serum blocking solution, washed, and mounted on slides with Mowiol 4-88 (48 mg/ml). In experiments using BW723C86, 5-HT, or 8-OH-DPAT, the drug was diluted in imaging medium and applied to neurons for 10 or 20 min at 1 µM final concentration before fixation and immunolabeling. The imaging medium consisted of phenol red-free MEM supplemented with glucose (33 mM), Hepes (20 mM), glutamine (2 mM), Na+-pyruvate (1 mM), and B27 (1X, Invitrogen). Sets of neurons to be compared were labeled and imaged simultaneously.

### Fluorescence image acquisition and cluster analyses

Fluorescence image acquisition and analyses were performed as previously described (Benhadda et al., 2021). Images were obtained on a Leica SP5 confocal microscope using the LAS-AF program (Leica). Stacks of 16–35 images were acquired using a 100 X objective with an interval of 0.2 µm and an optical zoom of 1.5. Image exposure time was determined on bright cells to avoid pixel saturation. All images from a given culture were then acquired with the same exposure time. Quantifications of 5-HT_2B_-R clusters were performed using MetaMorph software (Roper Scientific) on projections (sum of intensity) of confocal optical sections. For each cell, a region of interest (ROI) was chosen. For 5-HT_2B_-R cluster analysis, images were first flattened background filtered (kernel size, 3 × 3 × 2) to enhance cluster outlines, and a user-defined intensity threshold was applied to select clusters and avoid their coalescence. Thresholded clusters were binarized, and binarized regions were outlined and transferred onto raw data to determine the mean 5-HT_2B_-R cluster number, area and fluorescence intensity. The dendritic surface area of regions of interest was measured to determine the number of clusters per 10 µm². As the 5-HT_1A_-R presents an expression pattern in small clusters, the average fluorescent intensity per pixel was measured in a defined ROI. For each culture, we analyzed 7-12 dendrites per experimental condition. A total of about 10 neurons were analyzed per condition from three to five independent cultures. The experimenter was blind to the culture treatment.

### STORM imaging

STochastic Optical Reconstruction Microscopy (STORM) imaging was conducted on an inverted N-STORM Nikon Eclipse Ti microscope equipped with a 100× oil-immersion TIRF objective (NA 1.49) and an Andor iXon Ultra 897 EMCCD camera using 405 and 638 nm lasers from Coherent. Movies of 30,000 frames were acquired at frame rates of 50 Hz. The z position was maintained during acquisition by a Nikon Perfect Focus System and multicolor fluorescent microspheres (Tetraspeck, Invitrogen) were used as markers to register long-term acquisitions and correct for lateral drifts. Single-molecule localization and 2D image reconstruction was conducted as described in (Specht et al., 2013) by fitting the point-spread function of spatially separated fluorophores to a 2D Gaussian distribution. The surface of clusters and the densities of molecules per µm^2^ were measured in reconstructed 2D images through cluster segmentation based on detection densities. The threshold to define the border was set to 1000 detections/µm^2^. All pixels containing <2 detections were considered empty, and their intensity value was set to 0. The intensity of pixels with 2 detections was set to 1. The resulting binary image was analyzed with the function “regionprops” of Matlab to extract the surface area of each cluster identified by this function. Density was calculated as the total number of detections in the pixels belonging to a given cluster, divided by the area of the cluster.

### Electrophysiology

For patch clamp experiments, 260 µm-thick brain coronal slices were prepared from cWT-RCE mice (control) and cKO^5-HT^-RCE mice at 4 weeks (25-31 postnatal days) that express GFP only in Pet1-positive 5-HT neurons. The brain extraction and slicing was performed in ice-cold (0– 4°C) oxygenated (95% O_2_-5% CO_2_) solution containing 110 mM choline chloride, 2.5 mM KCl, 25 mM glucose, 25 mM NaHCO_3_, 1.25 mM NaH_2_PO_4_, 0.5 mM CaCl_2_, 7 mM MgCl_2_, 11.6 mM L-ascorbic acid, and 3.1 mM sodium pyruvate. Slices were then stored in artificial cerebro-spinal fluid (aCSF) containing 125 mM NaCl, 2.5 mM KCl, 25 mM glucose, 25 mM NaHCO_3_, 1.25 mM NaH_2_PO_4_, 2 mM CaCl_2_, and 1 mM MgCl_2_ (pH 7.2, maintained by continuous bubbling with 95% O_2_-5% CO_2_). Slices were incubated in aCSF at 32°C for 20 min and then at room temperature (20–25°C). Slices were transferred to the recording chamber where they were continuously superfused with oxygenated aCSF (30–32°C). To block synaptic transmission, we added to the aCSF 6-cyano-7-nitroquinoxaline-2,3-dione (CNQX; 10 µM, Hello Bio), D-2-amino-5-phosphonopentanoic acid (D-APV; 50 µM, Hello Bio) and SR95531 hydrobromide (GABAzine, 10 µM, Hello Bio). We also used (±) 8-hydroxy-2- (dipropylamino)tetralin hydrobromide (8-OH-DPAT; 30nM, SIGMA) and Apamin (20nM, Hello Bio) to activate 5-HT_1A_-Rs and block SK channels, respectively. Current clamp and voltage clamp recordings were performed with pipettes (5- to 7-MΩ resistance) prepared from borosilicate glass (BF150-86-10; Harvard Apparatus) using a DMZ pipette puller (Zeitz). Pipettes were filled with an intracellular solution containing 105 mM K-gluconate, 10 mM HEPES, 10 mM phosphocreatine-Na, 4 mM ATP-Na_2_, and 30 mM KCl (pH 7.25, adjusted with KOH). The input resistance (Rinput) was computed using a 10-mV hyperpolarizing step from a holding potential of −65 mV (50 ms). Current-clamp and voltage-clamp recordings were performed using an EPC-10 amplifier (HEKA Elektronik). Data acquisition was performed using Patchmaster software (Heka Elektronik). The liquid junction potential (−5 mV) was left uncorrected. Signals were sampled at 20 kHz and filtered at 4 kHz. Data analysis was performed using Igor Pro (Wavemetrics). Statistical analyses were performed using Prism (GraphPad). The normality of data distribution was tested using Shapiro–Wilk’s test. Unpaired two-tailed t-tests (for normally distributed datasets) or Mann–Whitney tests (for non-normally distributed datasets) were used for comparisons between two groups. For multiple comparisons, we used two-way ANOVA followed by Sidak’s test. Values of *p* < 0.05 were considered statistically significant.

## Materials availability

All newly created materials can be accessed by asking the authors.

## Supporting information

Full statistics

## ACKNOWLEDGEMENTS

We thank the *Cell* and *Tissue Imaging Facility* of the Institut du Fer à Moulin (namely Theano Eirinopoulou, Mythili Savariradjane), where all image acquisitions and analyses have been performed, Aude Muzerelle for help with stereotaxic viral injections, and the staff of the IFM animal facility (namely Baptiste Lecomte, Gaël Grannec, François Baudon, Anna-Sophia Mourenco, Emma Courteau and Eloise Marsan).

This work has been supported by grants from the Agence Nationale de la Recherche (ANR-17-CE16-0008, ANR-11-IDEX-0004-02), the Fondation pour la Recherche Médicale (Equipe FRM DEQ2014039529) and the Fédération pour la Recherche sur le Cerveau (FRC-2019-19F10).

## Author contributions

C.L., S.L., L.M., and A.R. designed the studies. A.B., C.L, A.R and L.M. wrote the manuscript, all authors revised it. A.B., I.M., X. M., and M.R. performed pharmacological studies, BRET experiments, stereotaxic and *in-vivo* experiments, immunofluorescence, image acquisition and analysis with the help of S.L., C.D., and C.L. performed and analyzed the electrophysiology experiments. A.B., S.L., L.M., and A.R. performed data analysis.

