## Supplementary material for "5-HT_1A_ and 5-HT_2B_ receptor interaction and co-clustering regulates serotonergic neuron excitability": Full statistics

**Fig2.** Pharmacology of cells expressing 5-HT$_{2B}$-Rs with or without 5-HT$_{1A}$-Rs
**Fig2A** Mesulergine competition by BW
Ki values Mann-Whitney unpaired t-test. P=0.4848
**Fig2B** 8-OH-DPAT competion by NAN
Ki values Mann-Whitney unpaired t-test. P=0.8857
**Fig2C** Inositolphospate formation upon BW stimulation
EC50 values Mann-Whitney unpaired t-test. P=0.2403

**Fig3**
**Fig3A** Radioligand binding from intact cells expressing 5-HT$_{2B}$-Rs with or without 5-HT$_{1A}$-Rs
Comparison between Bmax, unpaired t-test with Welch's correction p=0.037

**Fig3C** Surface biotinylation performed on COS-7 cells expressing 5-HT$_{1A}$-Rs and/or 5-HT$_{2B}$-Rs
Ratio of total vs surface expression for 5-HT2B receptor unpaired t-test t=2.34 df=8 p=0.0474
Ratio of total vs surface expression for 5-HT1A receptor unpaired t-test t=0.777 df=8 p=0.459

**Fig4**
**Fig4A**
Ratio of total vs surface expression for 5-HT2B receptor unpaired t-test $t_4$=4.481 p=0.011

**Fig5**
**Fig5A**
5-HT2B-R Surface clusters, BW723C86 stimulation

| | | |
|---|---|---|
| Number | | |
| Kruskal-Wallis test | P value | <0.0001 |
| Dunn's multiple comp | Mean rank diff. | Adj P Val |
| 2B t0 number vs. 2B t10 number | 42.89 | 0.0026 |
| 2B t0 number vs. 2B t20 number | 67.32 | <0.0001 |
| 2B t0 number vs. 2B+1A t0 number | 43.99 | 0.0014 |
| Area | | |
| Kruskal-Wallis test | P value | <0.0001 |
| 2B t0 area vs. 2B t10 area | -44.61 | 0.0016 |
| 2B t0 area vs. 2B t20 area | -57.76 | <0.0001 |
| 2B t0 area vs. 2B+1A t0 area | -45 | 0.0008 |
| Integrated Intensity | | |
| Kruskal-Wallis test | P value | <0.0001 |

| | | |
|---|---|---|
| 2B t0 Integrated Intensity vs. 2B t20 Integrated Intensity | -48.01 | 0.0007 |
| 2B t10 Integrated Intensity vs. 2B t20 Integrated Intensity | -35.77 | 0.0421 |

**Fig5B**

5-HT1A-R Surface expression, 8-OH-DPAT stimulation

| | | |
|---|---|---|
| Kruskal-Wallis test | P value | <0.0001 |

| Dunn's multiple comp | Mean rank diff. | Adjusted P Value |
|---|---|---|
| 1A t0 vs. 1A t10 | 44.25 | 0.0115 |
| 1A t0 vs. 1A+2B t0 | -39.87 | 0.0172 |
| 1A t0 vs. 1A+2B t10 | -46.74 | 0.0069 |

**Fig6C**

STORM 5-HT2B-R Surface clusters, BW723C86 stimulation

Number

| | | |
|---|---|---|
| Interaction | $F_{1, 70} = 0.9756$ | P=0.3267 |
| Treat | $F_{1, 70} = 11.15$ | P=0.0014 |
| Exp | $F_{1, 70} = 25.31$ | P<0.0001 |

| Tukey's multiple comp | Mean Diff | 95% CI | Adj P Val |
|---|---|---|---|
| t0:5-$HT_{2B}$-R vs. t0:5-$HT_{2B}$-R+5-$HT_{1A}$-R | 51.61 | 19.69 to 83.53 | 0.0004 |
| t0:5-$HT_{2B}$-R vs. t20:5-$HT_{2B}$-R | 37.1 | 3.095 to 71.1 | 0.0271 |
| t0:5-$HT_{2B}$-R vs. t20:5-$HT_{2B}$-R+5-$HT_{1A}$-R | 71.77 | 39.85 to 103.7 | <0.0001 |
| t0:5-$HT_{2B}$-R+5-$HT_{1A}$-R vs. t20:5-$HT_{2B}$-R | -14.51 | -46.43 to 17.41 | 0.6311 |
| t0:5-$HT_{2B}$-R+5-$HT_{1A}$-R vs. t20:5-$HT_{2B}$-R+5-$HT_{1A}$-R | 20.16 | -9.522 to 49.84 | 0.2879 |
| t20:5-$HT_{2B}$-R vs. t20:5-$HT_{2B}$-R+5-$HT_{1A}$-R | 34.67 | 2.754 to 66.59 | 0.0280 |

Area

| | | |
|---|---|---|
| Interaction | $F_{1, 70} = 0.0629$ | P=0.8027 |
| Treat | $F_{1, 70} = 7.067$ | P=0.0097 |
| Exp | $F_{1, 70} = 16.78$ | P=0.0001 |

| Tukey's multiple comp | Mean Diff | 95% CI of diff | Adj P Value |
|---|---|---|---|
| t0:5-$HT_{2B}$-R vs. t0:5-$HT_{2B}$-R+5-$HT_{1A}$-R | -191.7 | -377.2 to -6.161 | 0.0402 |
| t0:5-$HT_{2B}$-R vs. t20:5-$HT_{2B}$-R | -120 | -317.6 to 77.65 | 0.3865 |
| t0:5-$HT_{2B}$-R vs. t20:5-$HT_{2B}$-R+5-$HT_{1A}$-R | -336.7 | -522.2 to -151.2 | <0.0001 |
| t0:5-$HT_{2B}$-R+5-$HT_{1A}$-R vs. t20:5-$HT_{2B}$-R | 71.67 | -113.8 to 257.2 | 0.7402 |
| t0:5-$HT_{2B}$-R+5-$HT_{1A}$-R vs. t20:5-$HT_{2B}$-R+5-$HT_{1A}$-R | -145 | -317.5 to 27.52 | 0.1300 |
| t20:5-$HT_{2B}$-R vs. t20:5-$HT_{2B}$-R+5-$HT_{1A}$-R | -216.7 | -402.2 to -31.16 | 0.0156 |

**Fig7**

**Fig7B**

5-HT1A-R Surface expression. BW723C86 stimulation

One way ANOVA $F_{2. 43} = 16.15$ P value <0.0001

| Tukey's multiple comp | Mean diff. | Adjusted P Value |
|---|---|---|
| 5-HT1A-R t0 vs. 5-HT1A-R+5-HT2B-R t0 | -27.87 | 0.0188 |
| 5-HT1A-R t0 vs. 5-HT1A-R+5-HT2B-R t20" | 27.35 | 0.0463 |

| | | |
|---|---|---|
| 5-HT1A-R+5-HT2B-R t0 vs. 5-HT1A-R+5-HT2B-R t20" | 55.22 | <0.0001 |

**Fig7C**

5-HT2B-R Surface expression. 8-OH-DPAT stimulation

Number

| | | | | |
|---|---|---|---|---|
| Kruskal-Wallis test | P value | <0.0001 | | |
| Dunn's multiple comp | | | Mean rank diff. | Adjusted P Value |
| 2B+1A t0 nb vs. 2B+1A t10 nb | | | 25.33 | <0.0001 |
| 2B+1A t0 nb vs. 2B+1A t20 nb | | | 20.93 | 0.0001 |

Area

| | | | | |
|---|---|---|---|---|
| Kruskal-Wallis test | P value | <0.0001 | | |
| 2B+1A t0 area vs. 2B+1A t10 area | | | -27.52 | <0.0001 |
| 2B+1A t0 area vs. 2B+1A t20 area | | | -21.82 | <0.0001 |

Integrated intensity

| | | | | |
|---|---|---|---|---|
| Kruskal-Wallis test | | P value | <0.0001 | |
| 2B+1A t0 Integrated Intensity vs. 2B+1A t10 II | - | 27.11 | | <0.0001 |
| 2B+1A t0 II vs. 2B+1A t20 II | | -23.13 | | <0.0001 |

**Fig8**

**Fig8A**

5-HT2B-R Surface expression. 5-HT stimulation

Number

| | | | | |
|---|---|---|---|---|
| Kruskal-Wallis test | P value | <0.0001 | | |
| Dunn's multiple comp | | | Mean rank | Adjusted P Value |
| 2B t0 nbr vs. 2B+1A t0 nbr | | | 40,63 | <0.0001 |
| 2B t0 nbr vs. 2B+1A t20 nbr | | | 60,67 | <0.0001 |
| 2B t20 nbr vs. 2B+1A t20 nbr | | | 67,58 | <0.0001 |

Area

| | | | | |
|---|---|---|---|---|
| Kruskal-Wallis test | P value | <0.0001 | | |
| 2B t0 area vs. 2B+1A t0 area | | | -43,91 | <0.0001 |
| 2B t0 area vs. 2B+1A t20 area | | | -61,21 | <0.0001 |
| 2B t20 area vs. 2B+1A t20 area | | | -66,09 | <0.0001 |

| | | |
|---|---|---|
| Kruskal-Wallis test | P value | =0.2387 |

**Fig8B**

5-HT1A-R Surfaceexpression. 5-HT stimulation

| | | |
|---|---|---|
| Kruskal-Wallis test | P value | <0.0001 |

| Dunn's multiple comp | Mean rank | Adjusted P Value |
|---|---|---|
| 1A t0 vs. 1A t20 | 50.2 | 0.0003 |
| 1A t0 vs. 2B+1A t20 | 48.18 | 0.0014 |
| 2B+1A t0 vs. 2B+1A t20 | 76.51 | <0.0001 |
| 2B+1A t20 vs. 2B+1A t20+RS | -49.73 | 0.0023 |

**Fig9**

**Fig9A**

Firing rate cWT vs $cKO^{5\text{-}HT}$ Two way ANOVA-Repeated measure

| | | |
|---|---|---|
| Interaction | $F_{13, 663} = 0.2221$ | P=0.9983 |
| Depolarization | $F_{13, 663} = 400.3$ | P<0.0001 |
| Genotype | $F_{1, 51} = 0.0353$ | P=0.8517 |

**Fig9B**

Resting membrane potential values cWT vs $cKO^{5\text{-}HT}$ Mann-Whitney unpaired t-test. P=0.1820

Input resistance values cWT vs $cKO^{5\text{-}HT}$ Mann-Whitney unpaired t-test. P=0.8255

**Fig9C**

Firing rate cWT vs $cKO^{5\text{-}HT}$ in the presence of 8-OH-DPAT; Two way ANOVA-Repeated measure

| | | |
|---|---|---|
| Interaction | $F_{13, 806} = 8.011$ | P<0.0001 |
| Depolarization | $F_{13, 806} = 536.6$ | P<0.0001 |
| Genotype | $F_{1, 62} = 23.74$ | P<0.0001 |

| Sidak's multiple comp | Mean Diff | 95% CI | Adj P Val |
|---|---|---|---|
| 200pA | -6,567 | -12.74 to -0.3976 | 0.0274 |
| 250pA | -7,907 | -14.08 to -1.738 | 0.0028 |
| 300pA | -9,466 | -15.64 to -3.296 | 0.0001 |
| 350pA | -10,12 | -16.29 to -3.948 | <0.0001 |
| 400pA | -10,77 | -16.94 to -4.601 | <0.0001 |
| 450pA | -10,98 | -17.15 to -4.813 | <0.0001 |
| 500pA | -11,18 | -17.35 to -5.014 | <0.0001 |
| 550pA | -12,05 | -18.22 to -5.881 | <0.0001 |
| 600pA | -11,2 | -17.37 to -5.026 | <0.0001 |
| 650pA | -11,28 | -17.45 to -5.107 | <0.0001 |

**Fig9D**

Resting membrane potential values cWT vs $cKO^{5\text{-}HT}$ in the presence of 8-OH-DPAT
Mann-Whitney unpaired t-test. P=0.3588

Input resistance values cWT vs $cKO^{5\text{-}HT}$ in the presence of 8-OH-DPAT
Mann-Whitney unpaired t-test. P=0.6269

**Fig9E**

Firing rate cWT vs $cKO^{5\text{-}HT}$ in the presence of 8-OH-DPAT+Apamin; Two way ANOVA-RM

| | | |
|---|---|---|
| Interaction | $F_{9, 504} = 0.7975$ | P=0.6188 |
| Depolarization | $F_{9, 504} = 329$ | P<0.0001 |
| Genotype | $F_{1, 56} = 0.1706$ | P=0.6811 |

**Fig9F**

Resting membrane potential values in the presence of 8-OH-DPAT+Apamin
Mann-Whitney unpaired t-test. P=0.6668

Input resistance values in the presence of 8-OH-DPAT+Apamin
Mann-Whitney unpaired t-test. P=0.1196
